# Adaptive phototaxis of *Chlamydomonas* and the evolutionary transition to multicellularity in Volvocine green algae

**DOI:** 10.1101/2022.07.24.500296

**Authors:** Kyriacos C Leptos, Maurizio Chioccioli, Silvano Furlan, Adriana I Pesci, Raymond E Goldstein

**Author notes:** Pulmonary, Critical Care and Sleep Medicine, Department of Internal Medicine, Yale School of Medicine, 300 Cedar Street, New Haven, CT 06519, USA. Sensing Electromagnetic Plus Corp., 2450 Embarcadero Way, Palo Alto, CA-94303, USA.

## Abstract

A fundamental issue in biology is the nature of evolutionary transitions from unicellular to multicellular organisms. Volvocine algae are models for this transition, as they span from the unicellular biflagellate *Chlamydomonas* to multicellular species of *Volvox* with up to 50,000 *Chlamydomonas*-like cells on the surface of a spherical extracellular matrix. The mechanism of phototaxis in these species is of particular interest since they lack a nervous system and intercellular connections; steering is a consequence of the response of individual cells to light. Studies of *Volvox* and *Gonium*, a 16-cell organism with a plate-like structure, have shown that the flagellar response to changing illumination of the cellular photosensor is adaptive, with a recovery time tuned to the rotation period of the colony around its primary axis. Here, combining high-resolution studies of the flagellar photoresponse with 3D tracking of freely-swimming cells, we show that such tuning also underlies phototaxis of *Chlamydomonas*. A mathematical model is developed based on the rotations around an axis perpendicular to the flagellar beat plane that occur through the adaptive response to oscillating light levels as the organism spins. Exploiting a separation of time scales between the flagellar photoresponse and phototurning, we develop an equation of motion that accurately describes the observed photoalignment. In showing that the adaptive time scale is tuned to the organisms’ rotational period across three orders of magnitude in cell number, our results suggest a unified picture of phototaxis in green algae in which the asymmetry in torques that produce phototurns arise from the individual flagella of *Chlamydomonas*, the flagellated edges of *Gonium* and the flagellated hemispheres of *Volvox*.

## I. INTRODUCTION

A vast number of motile unicellular and multicellular eukaryotic microorganisms exhibits phototaxis, the ability to steer toward a light source, without possessing an image-forming optical system. From photosynthetic algae [1] that harvest light energy to support their metabolic activities to larvae of marine zooplankton [2] whose upward phototactic motion enhances their dispersal, the light sensor in such organisms is a single unit akin to one pixel of a CCD sensor or one rod cell in a retina [3]. In zooplanktonic larvae there is a single rhabdomeric photoreceptor cell [4] while motile photosynthetic microorganisms such as green algae [5] have a “light antenna” [6], which co-localizes with a cellular structure called the *eyespot*, a carotenoid-rich orange stigma. For these simple organisms, the process of *vectorial phototaxis*, motion in the direction of a source rather than in response to a light gradient [7], relies on an interplay between the detection of light by the photosensor and changes to the actuation of the apparatus that confers motility, namely their one or more flagella. Evolved independently many times [8], the common sensing/steering mechanism seen across species involves two key features.

The first attribute is a photosensor that has *directional* sensitivity, detecting only light incident from one side. It was hypothesized long ago [6] that in green algae this asymmetry could arise if the layers of carotenoid vesicles behind the actual photosensor act as an interference reflector. In zooplankton this “shading” role is filled by a single pigment cell [4]. This directionality hypothesis was verified in algae by experiments on mutants without the eyespot, that lacked the carotenoid vesicles [9], so that light could be detected whatever its direction. Whereas wild-type cells performed positive phototaxis (moving toward a light source), the mutants might naively have been expected to be incapable of phototaxis. Yet, they exhibited *negative* phototaxis, a fact that was explained as a consequence of an additional effect first proposed earlier [10]; the algal cell body functions as a convex lens with refractive index greater than that of water. Thus, a greater intensity of light falls on the photosensor when it was illuminated from behind than from the front, and a cell facing away from the light erroneously continues swimming in that direction, as if it were swimming toward the light.

The second common feature of phototactic microorganisms is a natural swimming trajectory that is helical. Spiral swimming has been remarked upon since at least the early 1900s, when Jennings [11] suggested that it served as a way of producing trajectories that are straight on the large scale, while compensating for inevitable asymmetries in the body shape or actuation of cilia, and Wildman [12] presciently observed that chirality of swimming and ciliary beating must ultimately be understood in terms of the genetic program contained within chromosomes. While neither offered a functional purpose related to phototaxis, Jennings did note earlier [13, 14] that when organisms swim along regular helices they always present the same side of their body to the outside. This implies that during regular motion the photosensor itself also has a fixed relationship to the helix.

In *Chlamydomonas*, motility derives from the breast-stroke beating of two oppositely-oriented flagella emanating from near the anterior pole of the cell body, as depicted in Fig. 1. The flagella, termed *cis* and *trans* for their proximity to the eyespot, define a plane, the unit normal to which is the vector **ê**_1_. Historical uncertainties around the precise three-dimensional swimming motion of *Chlamydomonas* were resolved with the work of Kamiya and Witman [15], the high-speed imaging study of Rüffer and Nultsch [16] and later work by Schaller *et al*. [17], who together demonstrated three features: (i) the eyespot is typically located on the equatorial plane of the cell, midway between **ê**_1_ and the vector **ê**_2_ that lies within the flagellar plane, pointing toward the *cis* flagellum, (ii) cells rotate counterclockwise (when viewed from behind) the axis **ê**_3_ at frequency *f*_r_ ∼ 1.5 − 2.5 Hz (*f*_*r*_ = 1.67±0.35 Hz in a recent direct measurement [18]), and (iii) positively phototactic cells swim along helices such that the eyespot always faces *outward*. The rotation around **ê**_3_ was conjectured to arise from a small non-planarity of the beat, as has been recently verified [19], while helical swimming arises from rotation around **ê**_1_ due to a slight asymmetry in the two flagellar beats.

**FIG. 1.**
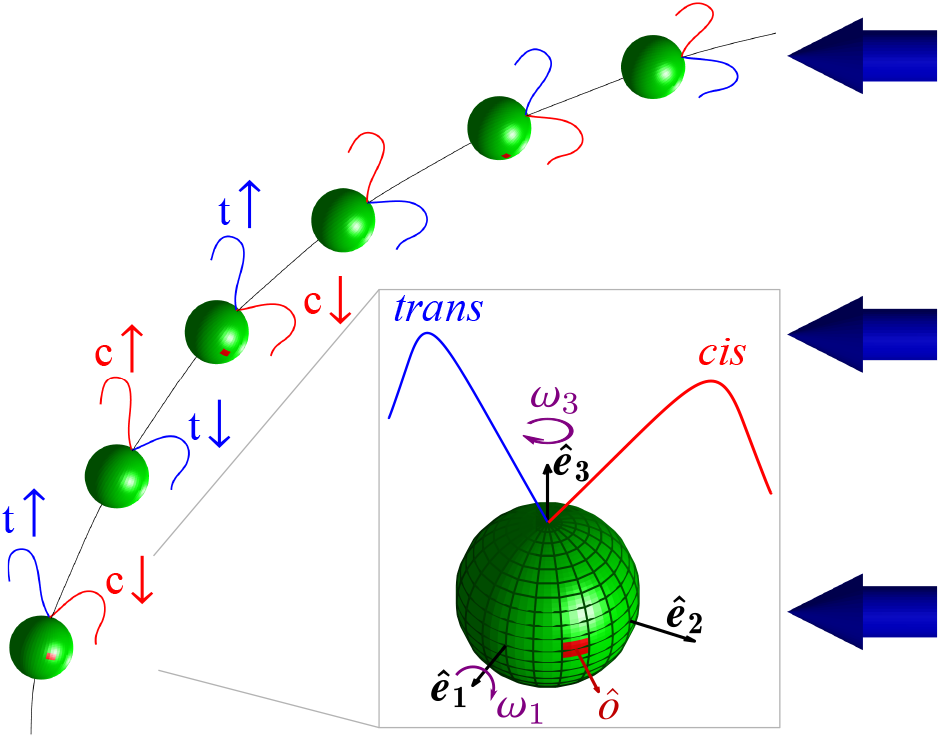
Phototaxis in *Chlamydomonas*. Cell geometry (in box) involves the primary axis **ê**_3_ around which the cell spins at angular frequency *ω*_3_, the axis **ê**_2_ in the flagellar beat plane, and **ê**_1_ = **ê**_2_ *×* **ê**_3_. As the cell swims and spins around **ê**_3_, its eyespot (red) moves in an out of the light shining in the direction of the blue arrows. This periodic light stimulation leads to alternating *cis* and *trans* flagella dominance, producing rotations around **ê**_**1**_ and hence a phototactic turn.

It follows from the above that the eyespot of a cell whose swimming is not aligned to the light receives an oscillating signal at angular frequency *ω*_3_ = 2*πf*_r_. Detailed investigation into the effect of this periodic signal began with the work of Rüffer and Nultsch, who used cells immobilized on micropipettes to enable high-speed cinematography of the waveforms. Their studies [20, 21] of beating dynamics in a negatively-phototactic strain showed the key result that the *cis* and *trans* flagella responded differently to changing light levels by altering their waveforms in response to the periodic steps-up and steps-down in signals that occur as the cell rotates. This result led to a model for phototaxis [17] that divides turning into two phases (Fig. 1): *phase I*, in which the eyespot moves from shade to light, causing the *trans* flagellum to increase transiently its amplitude relative to the *cis* flagellum, and *phase II*, in which the eyespot moves from light to shade, leading to transient beating with the opposite asymmetry. Both phases lead to rotations around **ê**_1_, and turns toward the light. The need for an asymmetric flagellar response was shown in studies of the mutant *ptx1* [22, 23], which lacks calcium dependent flagellar dominance [24] and can not do phototaxis.

These transient responses were studied further [25] through the photoreceptor current (PRC) that can be measured in the surrounding fluid. Subjecting a suspension of immotile cells (chosen to avoid movements) to rectified sinusoidal light signals that mimic those received by a rotating cell, they found that the PRC amplitude displays a maximum as a function of frequency, with a peak close to the body rotation frequency *f*_*r*_. This “tuning” of the response curve was investigated in more detail—in a negatively-phototactic strain—in the important work of Josef, et al. [26], who projected the image of the cell onto a quadrant photodiode whose analog signal could be digitized at up to 4000 samples per second. While this device did not allow detailed imaging of the entire waveform, it was able to capture changes in the forward reach of the two flagella (termed the “front amplitude”) over significantly longer time series than previous methods. Combined with later work that analyzed the signals within the framework of linear systems analysis [27], these studies showed how each of the two flagella exhibits a distinct, peaked frequency response.

From the original measurements of transient PRCs induced by step changes in light levels [25], it was evident that the response in time was biphasic and *adaptive* —a rapid rise in signal accompanied by slower recovery phase back to the resting state —and the presence of two timescales is implicit in the existence of the peak in the frequency response. More recently, measurements of the flagella-driven fluid flow around colonies of the multicellular alga *Volvox carteri* [28] showed again this adaptive response, which could be described quantitatively by a model previously to describe chemotaxis of both bacteria [29] and spermatozoa [30]. In a suitably rescaled set of units, the two variables *p* and *h* in this model respond to a signal *s*(*t*) through the coupled ODEs

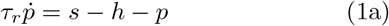

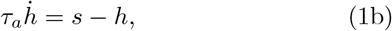

where *p* governs some observable, *h* represents hidden biochemistry responsible for adaptation, *τ*_*r*_ is the rapid response time, and *τ*_*a*_ is the slower adaption time. In bacteria, the adaptive response is exhibited by the biochemical network governing rotation of flagella, while for sperm curvature of the swimming path was altered linearly with *p* in response to a chemoattractant.

The model (1) was incorporated into a theory of *Volvox* phototaxis using a coarse-grained description of flagella-driven flows akin to the squirmer model [31], with a dynamic slip velocity **u**(*θ, ϕ, t*) as a function of spherical coordinates on the colony surface. Without light stimulation, the velocity is an axisymmetric function **u**_0_(*θ*) that varies with the polar angle *θ*, and is dominated by the first mode *u*_1_ ∝ sin *θ* [32]. As (1) is meant to describe the fluid flow associated with flagella of each of the somatic cells on the surface, it is introduced into the slip-velocity model through response *fields p*(*θ, ϕ, t*) and *h*(*θ, ϕ, t*) over the entire surface. Experiments indicate that the photoresponse is accurately represented the form

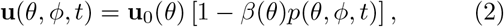

where the parameter *β* encodes the latitude-dependent photoresponse of the flagella (strong at the anterior of the colony, weak in its posterior). The swimming trajectories were then obtained from integral relationships between the slip velocity and the colony angular velocity [33].

Statistical analysis of many *Volvox* colonies shows that there is tuning of the response in that the product *f*_*r*_*τ*_*a*_ ≈ 1 (*f*_*r*_*τ*_*a*_ = 1.20 ± 0.44) [28], as indicated in Fig. 2. The significance of the product *f*_*r*_*τ*_*a*_ being of order unity can be understood as follows: when a region of somatic cells rotates to face a light source, the fluid flow it produces will decrease as *p* rapidly increases on a time scale *τ*_*r*_, and if the time *τ*_*a*_ it takes for *p* to recover is comparable to the colony rotation period 1*/f*_*r*_ then the fluid flow along the dark side will be stronger than than on the light side, and the colony turns to the light.

**FIG. 2.**
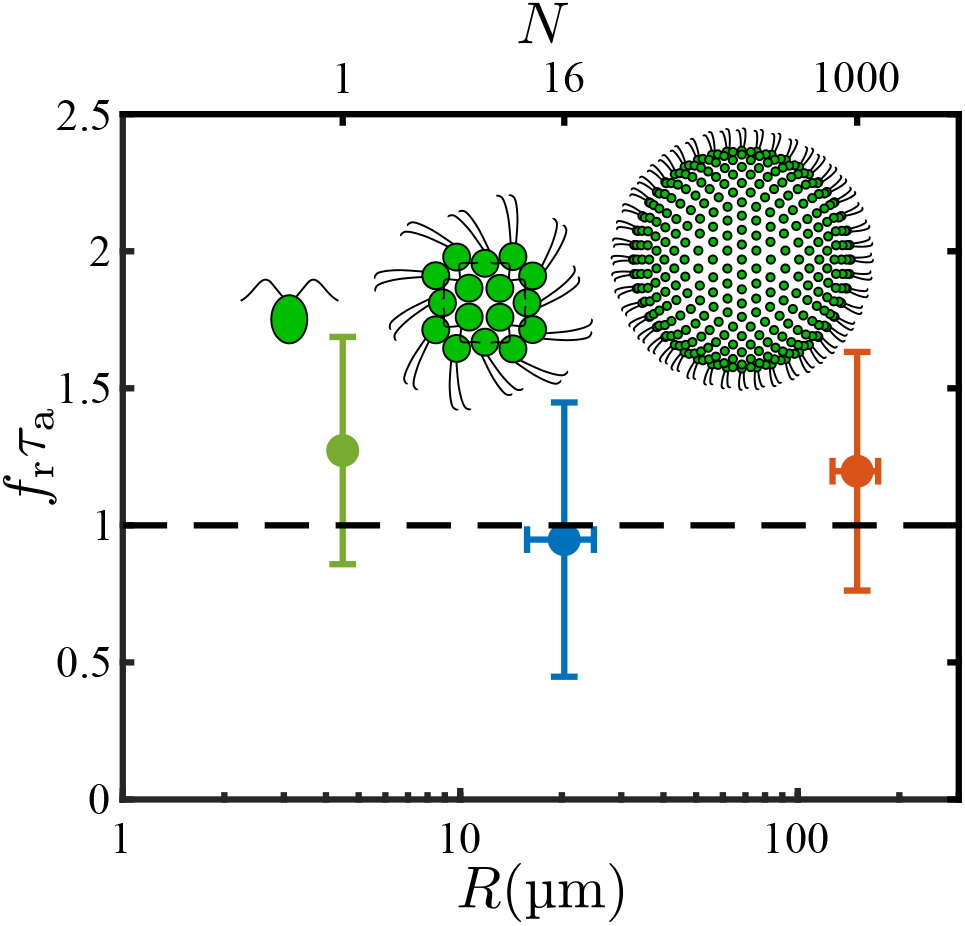
Master plot of adaptive time scales in Volvocine green algae. For each of *Chlamydomonas* (this paper), *Gonium* [34], and *Volvox* [28] the dimensionless product of the rotation frequency *f*_*r*_ around the primary body-fixed axis and the flagellar adaptive time *τ*_*a*_ is plotted as a function of the organism radius *R* (bottom axis) and typical cell number *N* (top axis).

A similar tuning phenomenon is found with *Gonium* [34], a member of the Volvocales typically composed of 16 cells arranged in a flat sheet as in Fig. 2. The flagella of the four central cells beat in a *Chlamydomonas*-like breaststroke waveform that propels the colony in the direction of the body-fixed axis **ê**_3_ perpendicular to the sheet. The flagella of the outer 12 cells beat at an angle with respect to the plane; their dominant in-plane component rotates the colony at frequency *f*_*r*_ about **ê**_3_, while the out-of-plane component adds to the propulsive force of the central cells. Experiments show that the peripheral cells display the same kind of biphasic, adaptive response as do *Volvox* colonies. This light-induced “drop-and-recover response” produces an axial force component *f*_‖_ from the peripheral flagella of the form

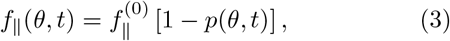

where 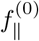 is the uniform component in the absence of photostimulation. Again, the directionality of the eyespot sensitivity leads to a photoresponse *p* that is greatest (and *f*_‖_ that is smallest) for those cells facing the light, and this nonuniformity in *f*_‖_ leads to a net torque about an in-plane axis which, balanced by rotational drag, leads to phototactic turning toward the light. The data for *Gonium* also supports tuning, with the product *f*_*r*_*τ*_*a*_ = 0.95 ± 0.50, as shown in Fig. 2.

In the present work we complete a triptych of studies in Volvocine algae by examining *Chlamydomonas*, the unicellular ancestor of all others [35]. Our purpose is to construct, in a manner that parallels that for *Volvox* and *Gonium*, a theory that links the photoresponse of flagella to the trajectories of cells turning to the light. We base the description on the kinematics of rigid bodies, where the central quantities are the angular velocities around body-fixed axes. This model bears some similarity to an earlier study of phototaxis [36], in which the asymmetric beating of flagella—modelled as spheres moving along orbits under the action of prescribed internal forces responding to light on the eyespot—was related to rotations about body-fixed axes, but the response to light was taken to be instantaneous and non-adaptive.

Results reported here on *Chlamydomonas* show that *f*_*r*_*τ*_*a*_ is close to unity (*f*_*r*_*τ*_*a*_ = 1.27 ± 0.41), from which we infer that tuning is an evolutionarily conserved feature spanning three orders of magnitude in cell number and nearly two orders of magnitude in organism radius (Fig. 2). We conclude that, in evolutionary transitions to multicellularity in the Volvocine algae, the ancestral photoresponse found in *Chlamydomonas* required little modification in order to work in vastly larger multicellular spheroids. The most significant change is basal body rotation [37] in the multicellulars in order that the two flagella on each somatic cell beat in parallel, rather than opposed as in *Chlamydomonas*. In *Gonium*, this arrangement in the peripheral cells leads to colony rotation, while for the somatic cells of *Volvox* the flagellar beat plane is tilted with respect to meridional lines, yielding rotation around the primary colony axis.

The presentation below proceeds from small scales to large, following a description in Sec. II of experimental methods used in our studies of the flagellar photoresponse of immobilized cells at high spatio-temporal resolution, and of methods for tracking phototactic cells. In Sec. III we arrive at an estimate of rotations about the body-fixed axis **ê**_1_ arising from transient flagellar asymmetries induced by light falling on the eyespot, and thus a protocol to convert measured flagella dynamics to angular velocities within the adaptive model. Section IV incorporates those results into a theory of phototactic turning. Exploiting a separation of time scales between individual flagella beats, cell rotation, and phototactic turning, we show how the continuous-time dynamics can be approximated by an iterated map, and allow direct comparison to three-dimensional trajectories of phototactic cells. By incorporating an adaptive dynamics at the microscale, one can examine the speed and stability of phototaxis as a function of the tuning parameter *f*_*r*_*τ*_*a*_ and deduce its optimum value. These results explain the many experimental results summarized above, and put on a mathematical basis phenomenological arguments [17] about the stability of phototactic alignment in *Chlamydomonas*.

## II. EXPERIMENTAL METHODS

### Culture conditions

Wild type *Chlamydomonas reinhardtii* cells (strain CC125 [38]) were grown axenically under photoautotrophic conditions in minimal media [39], at 23°C under a 100 μE·s^−1^ ·m^−2^ illumination in a diurnal growth chamber with a 14 : 10 h light-dark cycle.

### Flagellar photoresponse of immobilized cells

The flagellar photoresponse of *C. reinhardtii* was captured at high spatio-temporal resolution using the experimental setup shown in Fig. 3(a), which builds on previous studies [28, 40, 41]. Cells were prepared as described previously [41] – centrifuged, washed and gently pipetted into a bespoke observation chamber made of polydimethylsiloxane (PDMS). Chambers were mounted on a Nikon TE2000-U inverted microscope with a *×* 63 Plan-Apochromat water-immersion long-working-distance (LWD) objective lens (441470-9900; Carl Zeiss AG, Germany). Cells were immobilized via aspiration using a micropipette (B100-75-15; Sutter, USA) that was pulled to a *ø*5-μm tip, and the flagellar beat plane was aligned with the focal plane of the objective lens via a rotation stage. Video microscopy of immobilized cells was performed using a high speed camera (Phantom v341; Vision Research, USA) by acquiring 15 s movies at 2, 000 fps.

**FIG. 3.**
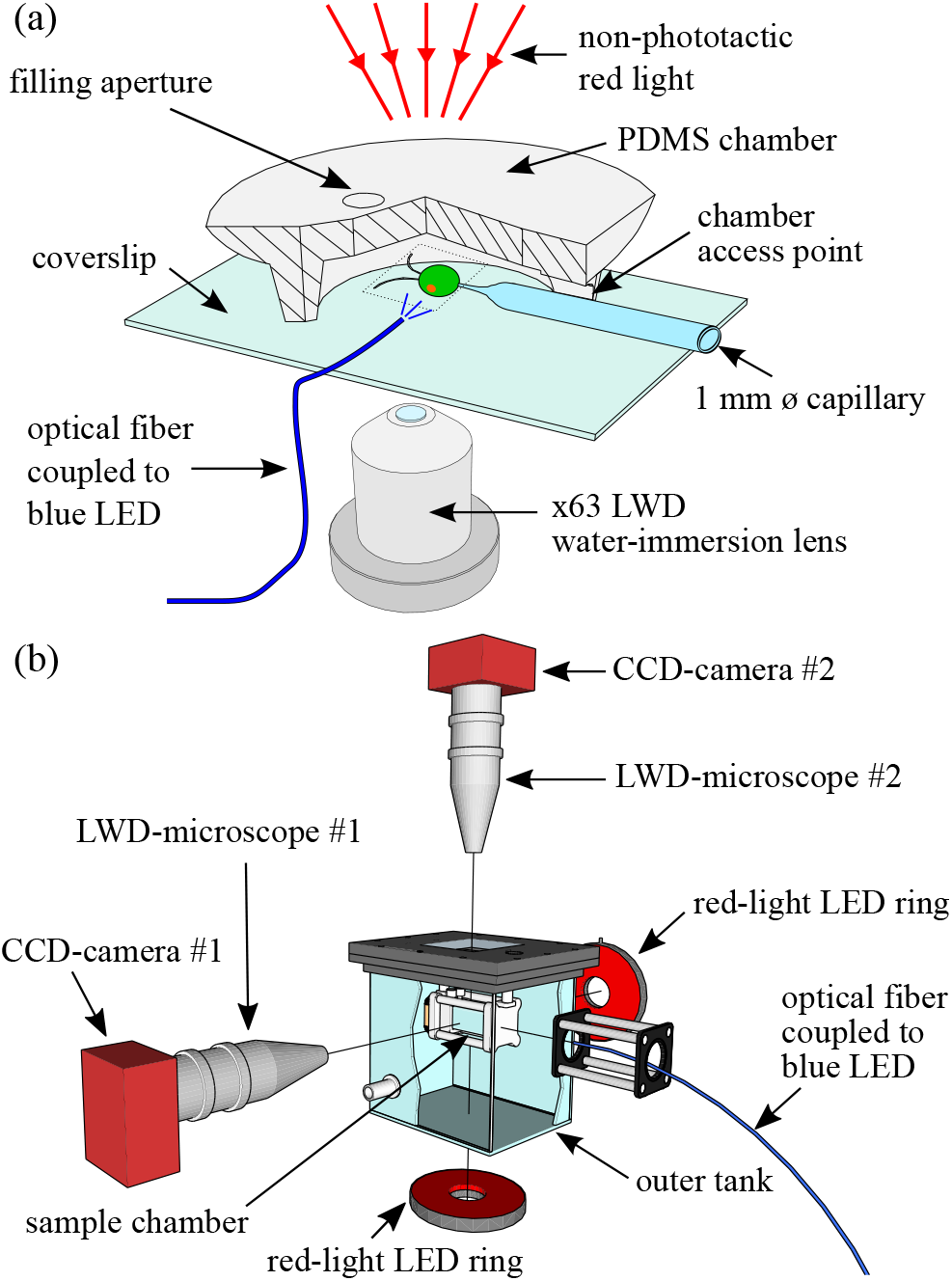
Experimental methods. Setups to measure (a) the flagellar photoresponse of cells immobilized on a micropipette, and (b) swimming trajectories of phototactic cells in a sample chamber immersed in an outer water tank to minimize thermal convection, as described in text.

The light used for photostimulation of cells was provided by a 470 nm Light Emitting Diode (LED) (M470L3; Thorlabs, USA) that was controlled via an LED driver (LEDD1B; Thorlabs, USA), coupled to a *ø*50 μm-core optical fiber (FG050LGA; Thorlabs, USA). This fiber is much smaller than that used in previous versions of this setup in order to accommodate the smaller size of a *Chlamydomonas* cell relative to a *Volvox* spheroid. The LED driver and the high-speed camera were triggered through a data-acquisition card (NI PCIe-6343; National Instruments, USA) using in-house programs written in `LabVIEW 2013` (National Instruments, USA), for both step- and frequency-response experiments. Calibration of the optical fiber was performed as follows: A photodiode (DET110; Thorlabs, USA) was used to measure the total radiant power *W* emerging from the end of the optical fiber for a range of voltage output values (0-5 V) of the LED driver. The two quantities were plotted and fitted to a power-law model which was close to linear.

Cells were stimulated at frame 2896 (≈1.45 s into the recording). A light intensity of ≈ 1 μE·s^−1^ ·m^−2^ (at 470 nm) was found empirically to give the best results in terms of reproducibility, sign, i.e. positive phototaxis, and quality of response; we conjecture that the cells could recover in time for the next round of stimulation. For the step-response experiments, biological replicates were *n*_cells_ = 3 with corresponding technical replicates *n*_tech_ = {4, 2, 2}. For the frequency-response experiments, biological replicates were *n*_cells_ = 3 with each cell stimulated to the following amplitude-varying frequencies: 0.5 Hz, 1 Hz, 2 Hz, 4 Hz and 8 Hz. Only the cells that showed a positive sign of response for *all* 5 frequencies are presented here. This was hence the most challenging aspect of the experimental procedure.

To summarize, the total number of high-speed movies acquired was *n*_mov[hs]_ = 24. All downstream analysis of the movies was carried out in `MATLAB`. Image processing and flagella tracking was based on previous work [41], and new code was written for force/torque calculations and flagellar photoresponse analysis.

### Phototaxis experiments on free-swimming cells

Three-dimensional tracking of phototactic cells was performed using the method described previously [42] with the modified apparatus shown in Fig. 3(b). The experimental setup comprised of a sample chamber suspended in an outer water tank to eliminate thermal convection. The modified sample chamber was composed of two acrylic flanges (machined in-house) that were clamped in a watertight manner onto an open-ended square borosilicate glass tube (2 cm *×* 2 cm *×* 2.5 cm; Vetrospec Ltd, UK). This design allowed a more accurate and easy calibration of the field of view and a simpler and better loading system of the sample via two barbed fittings. This new design also minimized sample contamination during experiments. Two 6 megapixel charge-coupled device (CCD) cameras (Prosilica GT2750; Allied Vision Technologies, Germany), coupled to two InfiniProbe™ TS160s (Infinity, USA) with Micro HM objectives were used to achieve a larger working distance than in earlier work (48 mm vs. 38 mm) at a higher total magnification of *×*16. The source of phototactic stimulus was a 470 nm blue-light LED (M470F1; Thorlabs, USA) coupled to a solarization-resistant optical fiber (M22L01; Thorlabs, USA) attached to an in-house assembled fiber collimator that included a *ø*12.7 mm plano-convex lens (LA1074A; Thorlabs, USA). Calibration of the collimated optical fiber was performed similarly to the experiments with immobilized cells. The calibration took account of the thickness of the walls of the outer water tank and the inner sample chamber, as well as the water in between.

The two CCD cameras and the blue-light LED used for the stimulus light were controlled using `LabVIEW 2013` (National Instruments, USA) including the image acquisition driver `NI-IMAQ` (National Instruments, USA). The cameras were triggered and synchronized at a frame rate of 10 Hz via a data-acquisition device (NI USB 6212-BNC; National Instruments, USA). For every tracking experiment (*n*_mov[3d]_ = 6), two 300-frame movies were acquired (side and top) with the phototactic light triggered at frame 50 (5 s into the recording). The intensity of the blue-light stimulus was chosen to be either 5 or 10 μE·s^−1^ ·m^−2^. To track the cells we used in-house tracking computer programs written in `MATLAB` as described in [42]. Briefly, for every pair of movies cells were tracked in the *side* and *top* movies corresponding to the *xz*-plane and in the *xy*-plane respectively. The two tracks were aligned based on their *x*-component to reconstruct the three-dimensional trajectories. The angle between the cell’s directional vector and the light was then calculated for every time point.

## III. FLAGELLAR DYNAMICS

### A. Forces and torques

We begin by examining the response of the two flagella of an immobilized *Chlamydomonas* cell to a change in the light level illuminating the eyespot. Figure 4 and Supplementary Video 1 [43] shows a comparison between the unstimulated beating of the flagella and the response to a simple step up from zero illumination. These are presented as overlaid flagellar waveforms during a the single beat in the dark and one that started 50 ms after the step. In agreement with previous work cited in Sec. I [21, 26], we see that the transient response involves the *trans* flagellum reaching further forward toward the anterior of the cell, while the *cis* waveform contracts dramatically. The photoresponse is adaptive; the marked asymmetry between the *cis* and *trans* waveforms decays away over ∼ 1 − 2 s, restoring the approximate symmetry between the two. This adaptive timescale is much longer than the period *T*_b_ ∼ 20 ms of individual flagellar beats.

**FIG. 4.**
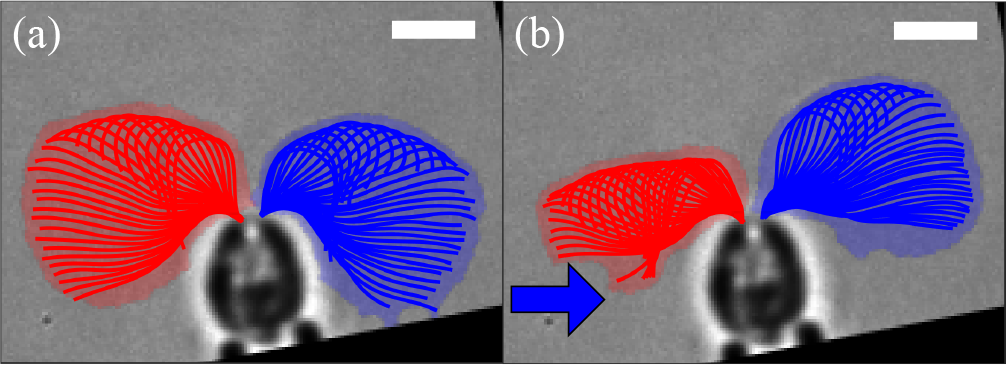
Flagellar photoresponse of an immobilized cell after a step-up in light. Light is from the left (blue arrow), toward the eyespot. The panels show overlaid flagellar waveforms of a single beat (a) in the dark and (b) starting at 50 ms (52.5 ms) for the *cis* (*trans*) flagellum after the step increase in light. Scale bar is 5 μm.

We wish to relate transient flagellar asymmetries observed with immobilized cells, subject to time-dependent light stimulation, to cell rotations that would occur for freely-swimming cells. We begin by examining the beating of unstimulated cells to provide benchmark observations. We analyze high-speed videos to obtain the waveforms of flagella of length *L*, radius *a*, in the form of moving curves **r**(*λ, t*) parameterized by arclength *λ* ∈ [0, *L*] and time. Within Resistive Force Theory (RFT) [44, 45], and specializing to planar curves, the hydrodynamic force density on the filament is

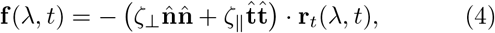

where 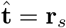 and 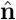 (with 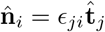) are the unit normal and tangent at *λ*, and *ζ*_⊥_ and *ζ*_‖_ are drag coefficients for motion perpendicular and parallel to the filament. We assume the asymptotic results *ζ*_⊥_ = 4*πμ/c*_⊥_ and *ζ*_‖_ = 2*πμ/c*_‖_, where 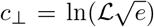 and 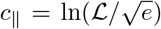, with *ℒ* = *L/a* the aspect ratio. Table I gives typical values of the cell parameters; with *ℒ* ≈ 108, we have *c*_⊥_ ≈ 5.2 and *c*_‖_ ≈ 4.2. To complete the analysis, we adopt the convention shown in Fig. 5(a) to define the start of a beat, in which chords drawn from the base to a point of fixed length on each flagellum define angles Θ_cis,trans_ with respect to **ê**_3_, Local minima in Θ_cis,trans_(*t*) [Fig. 5(b)] define the beat endpoints.

**TABLE I.** Geometry of flagellar beats. Data are from the present study except for the flagellum radius.

| Quantity | Symbol | Mean ± SD |
| --- | --- | --- |
| flagellum length | $L$ | $13.5 \pm 0.8 \mu\text{m}$ |
| flagellum radius [46] | $a$ | $0.125 \mu\text{m}$ |
| cell body radius | $R$ | $4.4 \pm 0.3 \mu\text{m}$ |
| beat frequency | $\hat{f}_b$ | $43.2 \pm 8.0 \text{ Hz}$ |
| anchor angle | $\varphi_a$ | $(0.05 \pm 0.01)\pi$ |
| initial angle | $\hat{\varphi}_0$ | $(0.26 \pm 0.05)\pi$ |
| sweep angle | $\hat{\varphi}_b$ | $(0.33 \pm 0.06)\pi$ |

**FIG. 5.**
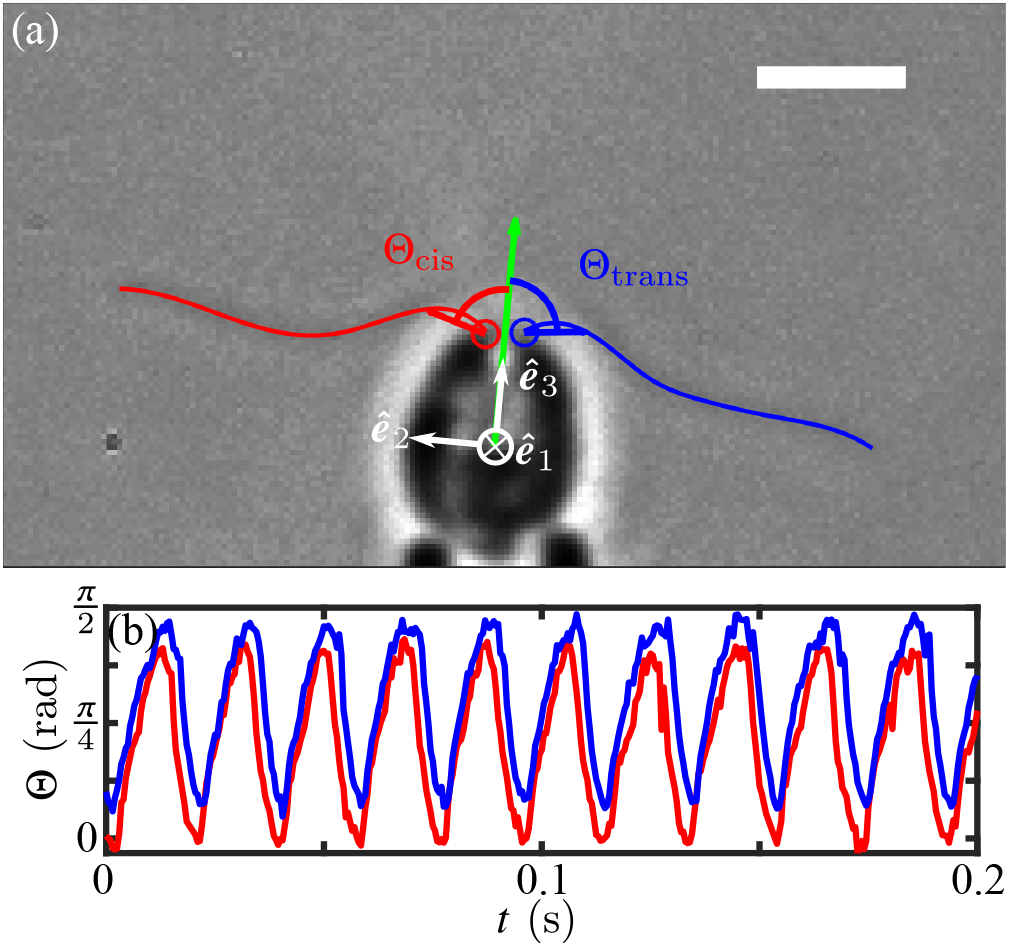
Flagellar beat cycles. (a) Angles Θ on each flagellum (red for *cis*, blue for *trans*) relative to symmetry axis **ê**_3_ (green) of the cell are used to define the cycle. Scale bar is 5 μm. (b) Typical time series of the two angles.

Using a hat (^) to denote quantities measured without photostimulation, Fig. 6 shows the results of this analysis for the propulsive component of the total force,

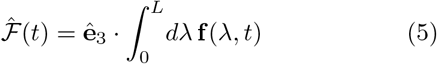

and the torque component around **ê**_1_,

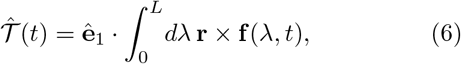

where **r** is measured from the cell center. The smoothness of the data arises from the large number of beats over which the data are averaged. The force 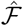 varies sinusoidally in time, offset from zero due to the dominance of the power stroke over the recovery stroke, with a peak value 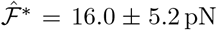 and mean over a beat period of 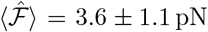 per flagellum. These findings are in general agreement with previous studies of *Chlamydomonas* cells held by an optical trap [47], micropipette-held cells [48], measurements on swimming cells confined in thin fluid films [49] and in bulk [50], and more recent work using using micropipette force sensors [51].

**FIG. 6.**
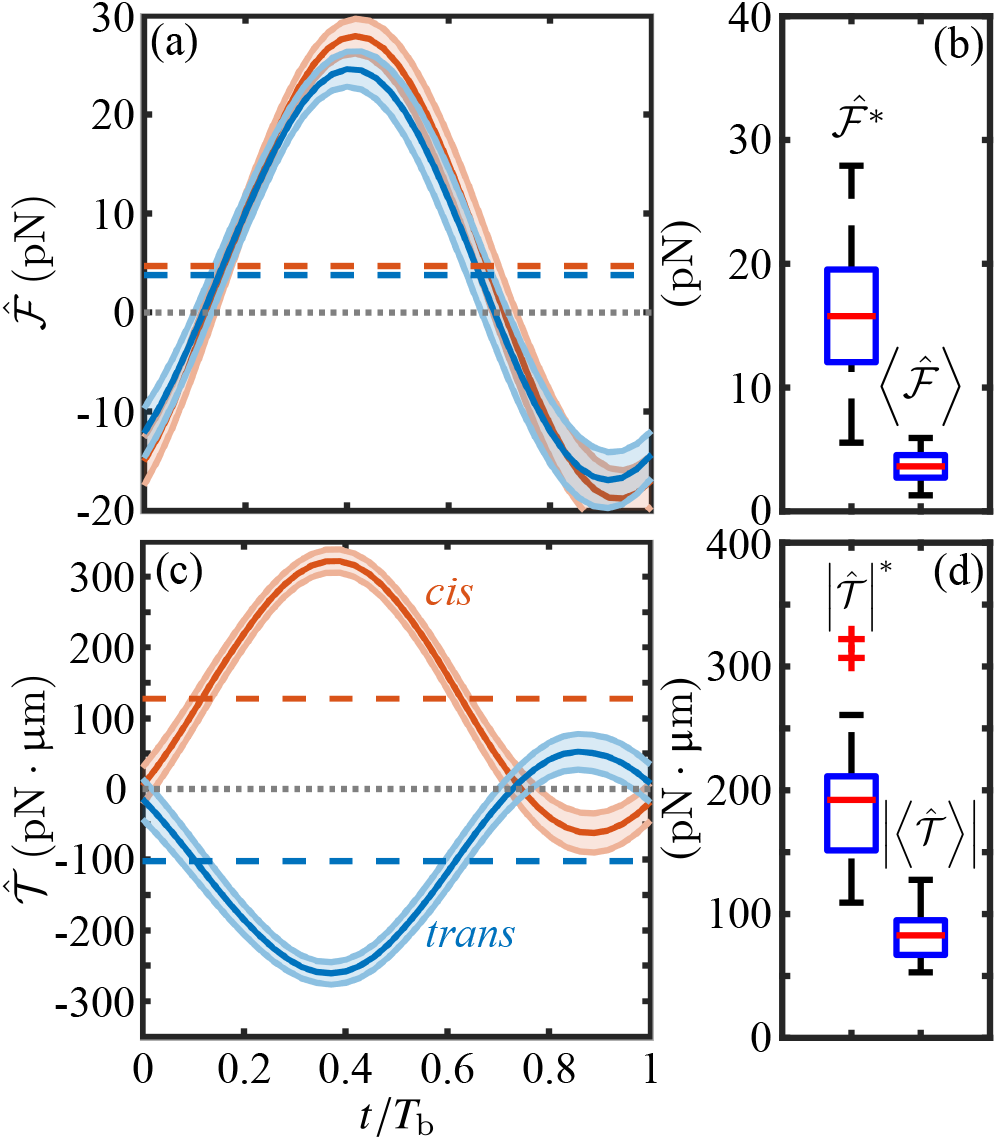
Flagellar forces and torques of unstimulated cells. Propulsive force (a) and torque about the cell center (c) of *cis* (red) and *trans* (blue) flagella during beat cycle of a representative cell, with cycle averages indicated by dashed lines. (b) and (d) show boxplots of peak values (^*^) and beat-average quantities (⟨ ⟩) computed from *n* = 48 flagella in 24 movies.

Aggregating all data obtained on the unstimulated torques exhibited by *cis* and *trans* flagella, we find a peak magnitude 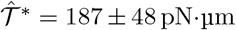 and cycle-average mean value 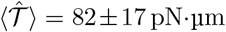. As a consistency check we note that the ratio torque/force should be an interpretable length, and we find 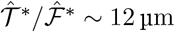, a value that is very close to the mean flagellar length *L* = 13.5 μm. Across a sample size of *n* = 24 we find the sum

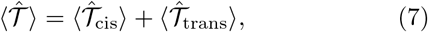

is 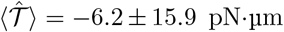, and thus is consistent with symmetry of the two flagella, but the data clusters into two clear groups; a *cis*-dominant subpopulation (*n* = 9), with 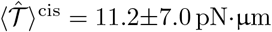 and a *trans*-dominant sub-population (*n* = 15) with 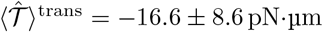. As discussed below, such differences would generally distinguish between positively and negatively phototactic cells, and the presence of both in our cellular population likely reflects the detailed growth and acclimatization conditions. For consistency, we focus here only on positively phototactic cells. For them, the residual torque provides information on the pitch and amplitude of helical trajectories of unstimulated cells.

### B. Heuristic model of flagellar beating

Below we extend the quantification of flagellar beating to a transient photoresponse like that in Fig. 4, with the goal of inferring the angular velocity *ω*_1_ around **ê**_1_ that a freely-swimming cell would experience, and which leads to a phototurn. The constant of proportionality between the torque and the angular velocity is an effective rotational drag coefficient that can be viewed as one of a small number of parameters of the overall phototaxis problem. Our immediate goal is to develop an estimate of this constant to provide a consistency check on the final theory in its comparison to experiment.

A simple model of the power stroke shown in Fig. 7 can be used to understand the peak values 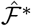 and 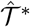: a straight flagellum attached at angle *φ*_*a*_ to a spherical body of radius *R*, whose beat angle *φ*(*t*) sweeps from 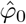 to 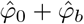. Table I and Fig. 8 summarize data on these geometric quantities. Relative to the center C a point at arclength *λ* ∈ [0, *L*] is at position

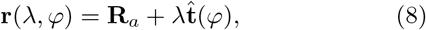

where **R**_*a*_ = *R* [sin *φ*_*a*_**ê**_*x*_ + cos *φ*_*a*_**ê**_*y*_] is the vector CO and 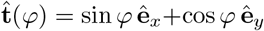 is the unit tangent. The velocity of a point on the filament is 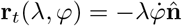.

**FIG. 7.**
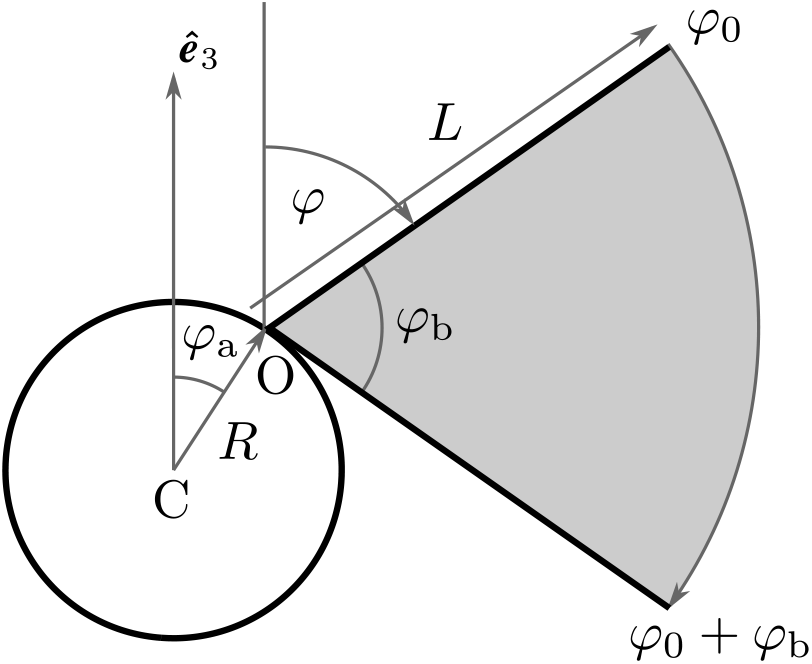
The pivoting-rod model of the power stroke.

**FIG. 8.**
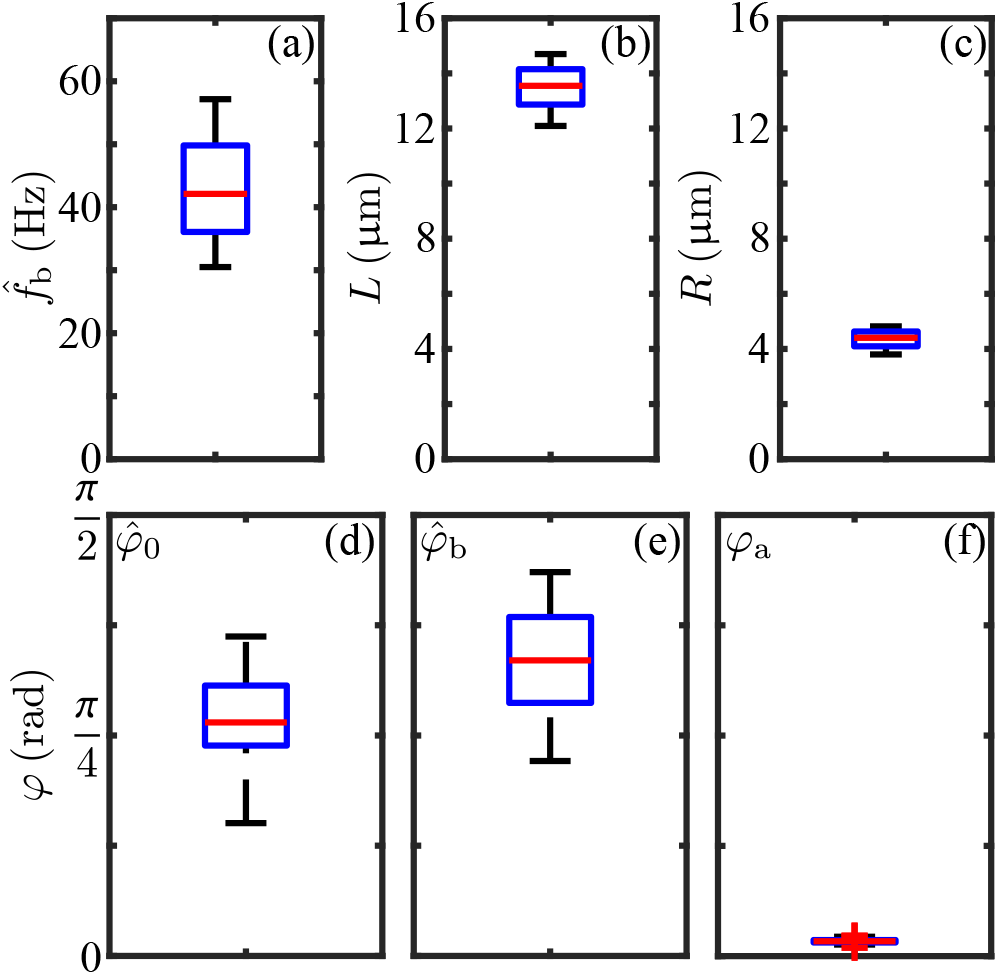
Distributions of geometric quantities for flagellar beats, computed from *n* = 24 movies: (a) instantaneous flagellar beat frequency 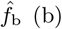 flagellar length *L*, (c) cell-body radius *R*, (d-f) per-beat flagella-line initial angle 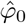, sweep angle 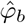 and anchor angle *φ*_*a*_.

The integrated force of the pivoting rod in Fig. 5 is

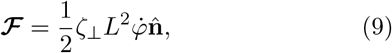

Considering the sinusoidal variation of the quantities in Figs. 6(a,c), we estimate *φ* by the lowest mode that has vanishing speed at beginning and end of the power stroke,

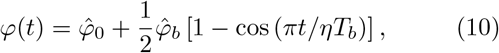

for *t* ∈ [0, *ηT*_*b*_], where *T*_*b*_ is the full beat period and *η* ≃ 0.7 is the fraction of the period occupied by the power stroke. Using the data in Table I, and the fact that the maximum projected force occurs very close to the time when *φ* = *π/*2, we obtain the estimate

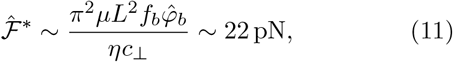

which is ∼ 1 s.d. above the experimental mean. While it is not surprising that the pivoting rod model overestimates the propulsive force relative to the actual undulating flagellum, the fact that this overestimate is small indicates that the essential physics is contained in (11).

Further heuristic insight into the flagellar forces produced can be gained by estimating the resultant motion of the cell body, assumed to be a sphere of radius *R*. This requires incorporating the drag of the body and that due to the flagella themselves. A full treatment of this problem requires going beyond RFT to account for the effect of flows due to the moving body on the flagellar and vice versa. In the spirit of the rod model, considerable insight can be gained in the limit of very long flagella, where the fluid flow is just a uniform translational velocity **u**(*t*) and the velocity of a point on the rod is

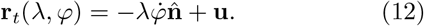

Symmetry dictates that the net force from the downward sweeps of two mirror-image flagella is along **ê**_3_, as is the translational velocity **u** = *u***ê**_*y*_ of the cell body. Adding mirror-image copies of the force (9) and the drag force on the body − *ζu***ê**_*y*_, where *ζ* = 6*πμR* is the Stokes drag coefficient for a sphere of radius *R*, the condition that the total force vanish yields

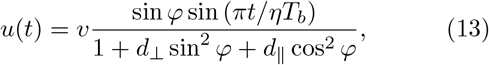

where *d*_⊥_ = 2*ζ*_⊥_*L/ζ* = 4*L/*3*Rc*_⊥_ and *d*_‖_ = 2*L/*3*Rc*_‖_. The speed *v* is given by the maximum force (11) as

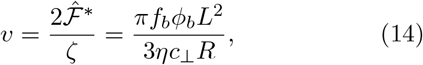

and is independent of the viscosity μ, as it arises from a balance between the two drag-induced forces of flagellar propulsion and drag on the spherical body, the latter ignoring the contribution in the denominator of (13) from flagellar drag. For typical parameters, *d*_⊥_ ≈ 0.8, and the denominator is ≈ 1.8 when *u* is maximized (at *φ* = *π/*2), while *υ* ∼ 540 μm/s. Thus, the peak swimming speed during the power stroke would be *u*^*^ *υ/*1.8 ∼ 300 μm/s, consistent with measurements [49], which also show that over a complete cycle, including the recovery stroke, the mean speed *ū* ∼ *u*^*^*/*4. We infer that *ū* ∼ 75 μm/s, consistent with observations [49, 52].

We now use the rod model to estimate the maximum torque 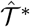 produced on an immobilized cell, to compare with the RFT calculation from the experimental waveforms. As in (12), the force density on the moving filament is 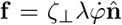, the torque density is 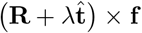, and the integrated torque component along **ê**_1_ is

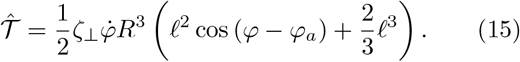

where 𝓁 = *L/R* and *φ* is again given by (10). The two terms in (15), scaling as *RL*^2^ and *L*^3^, arise from the distance offset from the cell body and the integration along the flagellar length, respectively.

Examining this function numerically we find that its peak occurs approximately midway through the power stroke, where *φ* − *φ*_*a*_ ≃ *π/*3, leading to the estimate

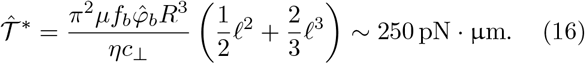

and, for average torque,

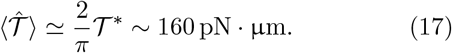

Here again these estimates are slightly more than 1 s.d. above the experimental value, giving further evidence that the rod model is a useful device to understand the scale of forces and torques of beating flagella.

The essential feature of (13) is an effective translational drag coefficient 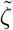 that is larger than that of the sphere due to the presence of the very beating flagella that cause the motion. For flagella oriented at 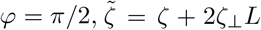, a form that reflects the extra contribution from transverse drag on the two flagella. We now consider the analogous rotational problem and estimate an effective rotational drag coefficient 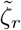 in terms of the bare rotational drag *ζ*_*r*_ = 8*πμR*^3^ for a sphere. If we set in rotational motion at angular speed Ω a sphere with two flagella attached at angles ±*φ*_*a*_, the velocity of a rod segment at *λ* is **Ω** *×* **r** and the calculation of the hydrodynamic force and torque proceeds as before, yielding

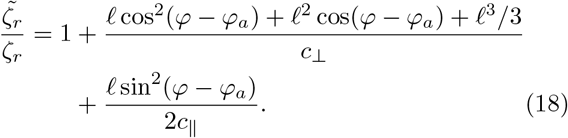

For typical parameters, *ζ*_*r*_ ∼ 2.1 pN·μm·s, and the added drag of the flagella is significant; the ratio in (18) varies from 4.8− 3 as *φ* varies from 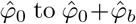. At the approximate peak of the power stroke we find 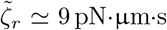, a value we use in further estimates below.

The effective rotational drag coefficient can be used to estimate the unstimulated angular speed due to the small torque imbalance noted below (7) for the *cis*-dominant subpopulation,

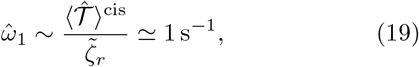

which can be compared to the angular speed |*ω*_3_| ≃ 10 s^−1^ of spinning around the primary axis. In Sec. IV we show that the small ratio 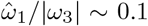 implies that the helices are nearly straight.

### C. Adaptive dynamics

The results of the previous section constitute a quantitative understanding of the phototactic torques produced *within* a given flagellar beat, which typically lasts 20 − 25 ms. As mentioned previously, the timescale for the full photoresponse associated with a change in light levels falling on the eyespot is considerably longer, on the order of 0.5 s. This separation of timescales is illustrated in Fig. 9, where we have schematically shown the time-resolved, oscillating phototactic torque of each of the two flagella, the signed sum, and its running average. It is precisely because of the separation of timescales between the rapid beating and both the slow response and the slow phototurns that a theory developed in terms of the beat-averaged torques is justified.

**FIG. 9.**
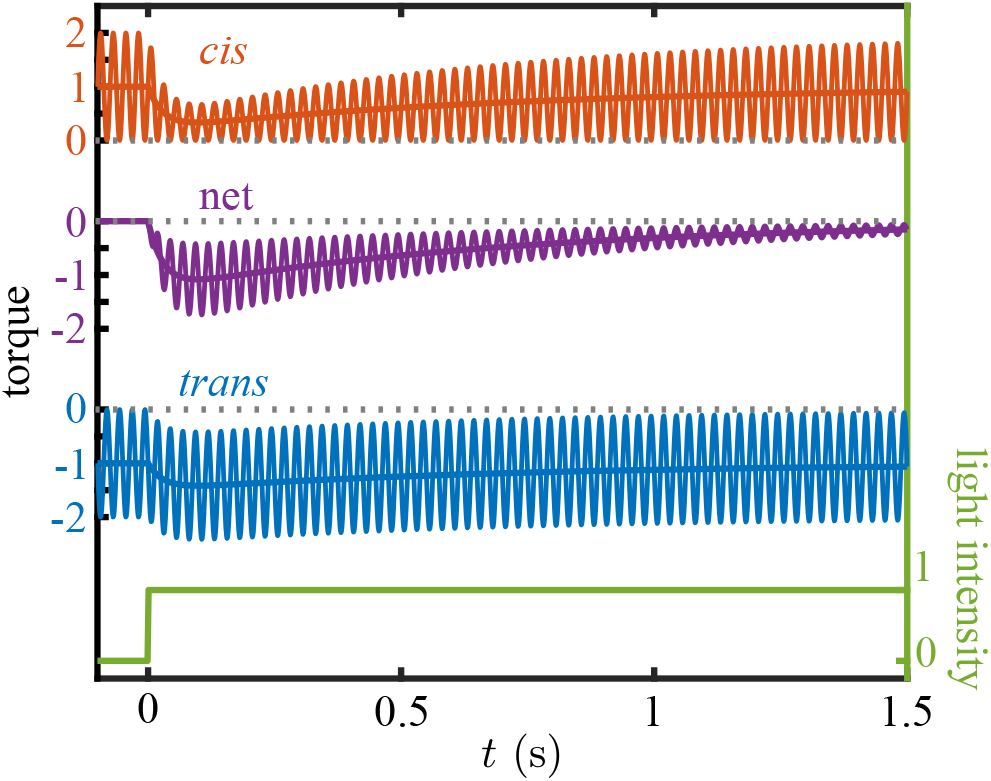
Schematic of flagellar photoresponse. A step-up in light at *t* = 0 (green) leads to a biphasic decrease in the mean value and oscillation amplitude of the *cis* phototorque (red), and a biphasic increase in the magnitude of the mean value of the *trans* phototorque (blue). The net torque (purple), the signed sum of the two contributions, has a biphasic response in both the oscillation amplitude and its running mean value.

In the following, we measure phototorques relative to the unstimulated state of the cell, and define the two (signed) *beat-averaged* quantities

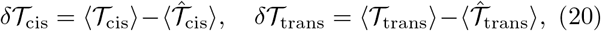

and their sum, the net beat-averaged phototactic torque

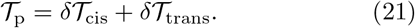

𝒯_p_ > 0 when the *cis* flagellum beats more strongly and 𝒯_p_ < 0 when the *trans* flagellum does. Our strategy is to determine 𝒯_p_ from experiment on pipette-held cells and to estimate the resulting angular speed 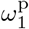 using 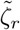.

The scale of net torques expected during a transient photoresponse can be estimated from the pivoting-rod model. From step-up experiments such as that shown in Fig. 4, we observe that there are two sweep angles 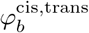 whose difference 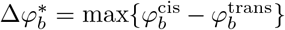 can be used in (17) to obtain 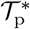, the maximum value of the beat-averaged sum (corresponding to the most negative value of the purple running mean in Fig. 9). Averaging over 8 photoresponse videos, we find 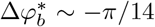, which yields the estimate

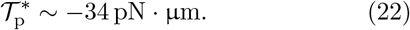

From the effective drag coefficient the corresponding peak angular speed in such a photoresponse is

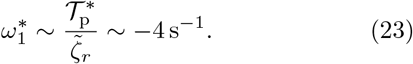

To put this in perspective, consider the photoalignment of an alga swimming initially perpendicular to a light source. If sustained continuously, complete alignment would occur in a time 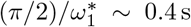, whereas our observations suggest a longer timescale of ∼ 2 s. This will be shown to follow from the variability of *ω*_1_ during the trajectory in accord with an adaptive dynamics.

While the estimate in (22) gives a guide to scale of the torques responsible for phototurns, we may calculate them directly within RFT from flagellar beating asymmetries in the same manner as in the unstimulated case. Figure 10 shows the response of a single cell to a step-up in illumination (of which Fig. 4 is a snapshot), in which the results are presented both in terms of 𝒯_p_ and the estimated 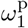. To obtain these data, the oscillating time series of *cis* and *trans* torques were processed to obtain beat-averaged values whose sum yields the running average, as in Fig. 9. The overall response is < 0, indicating that the *trans* flagellum dominates, and the peak value averaged over multiple cells of −37±12 pN·nm is consistent with the estimate in (22). The biphasic response, with a rapid increase followed by a slow return to zero, is the same form observed in *Volvox* [28] and *Gonium* [34]. We now argue that the adaptive model (1) used for those cases can be recast as an evolution equation for the angular speed itself, setting *p* = *ω*_1_ and *s* = *gI*, with *I* the light intensity and *g* a proportionality constant,

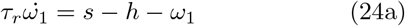

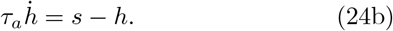

For constant *s* the system (24) has the fixed point 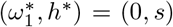. If *ω*_1_ = *h* = *s* = 0 for *t* ≤ 0, followed by *s* = *s*_0_ for *t* > 0, then

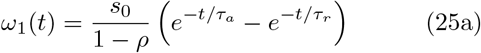

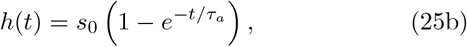

where *ρ* = *τ*_*r*_*/τ*_*a*_. The result (25a), illustrated in Fig. 11(a) for the case *ρ* = 0.1 and a square pulse of duration long compared to *τ*_*a*_, shows clearly the biphasic response of the data in Fig. 10. This behavior is like two coupled capacitors charging and discharging one another, particularly in the limit *ρ* ≪ 1. At early times, *h* remains small and *ω*_1_ relaxes toward *s*_0_ with the rapid timescale *τ*_*r*_. Later, when *t* ∼ *τ*_*a*_, *h* relaxes toward *s*_0_, and *ω*_1_ relaxes instead toward zero, completing the pulse.

**FIG. 10.**
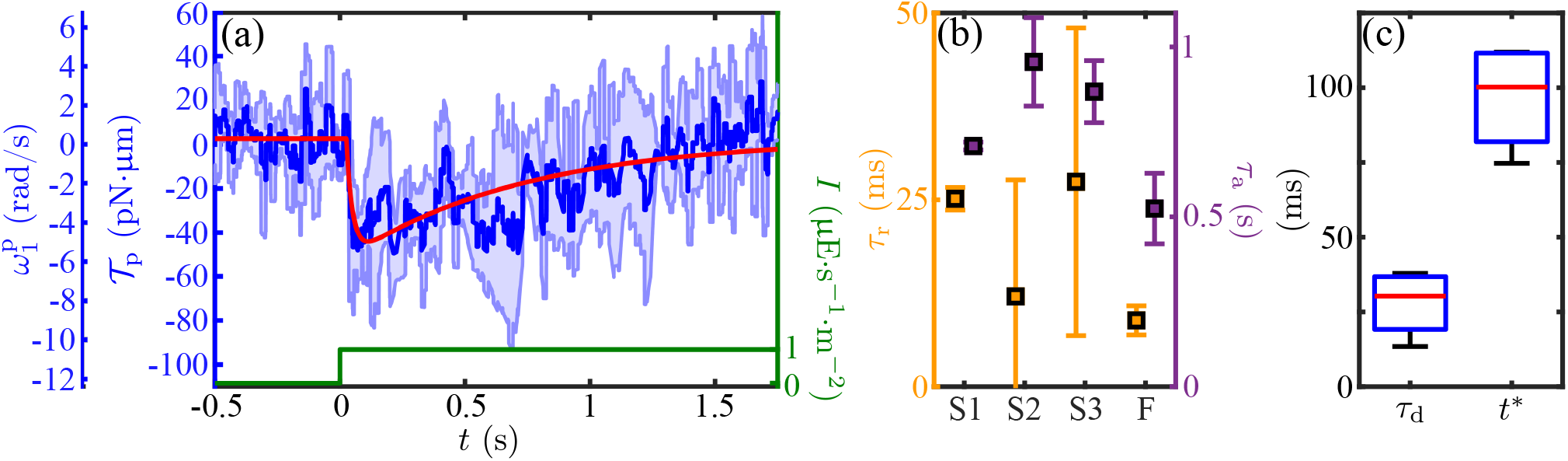
Dynamics of the flagellar photoresponse. (a) The mean (dark blue) and standard deviation (light-blue) of the proxy torque during a step-up stimulus for one cell (*n*_tech_ = 4) fitted to (25a) (red line). (b) Inset showing fitted (*τ*_*r*_, *τ*_*a*_) pairs for *n*_cells_ = 4 upon step-up stimulation. (c) As in (b) but for the delay time *τ*_*d*_ and time of maximum photoresponse amplitude *t*^*^.

**FIG. 11.**
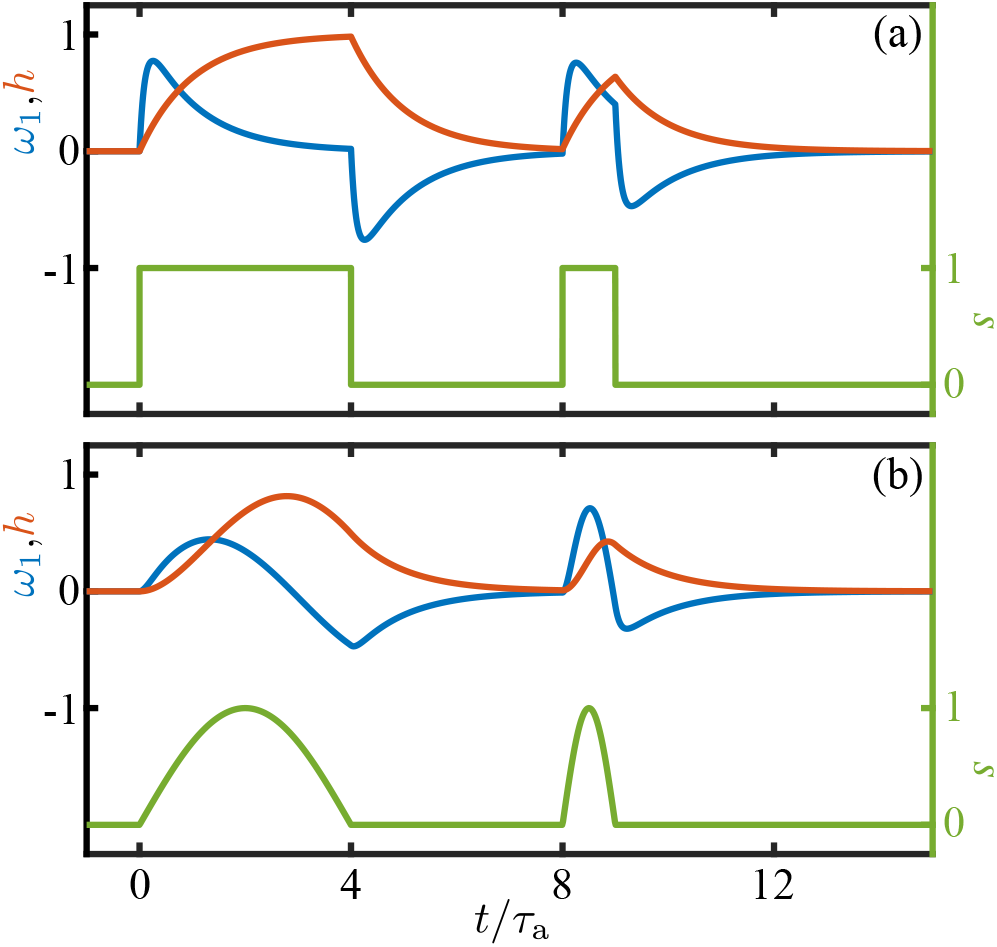
Dynamics of the adaptive model. (a) Response of the variables *p* and *h* to a square pulse of stimulus (green), for *ρ* = 0.1. (b) Response to rectified sinusoids.

After a step up, *h* has relaxed to −*s*_0_, and if *s* is then stepped down to zero, *ω*_1_ rapidly tends toward *s*_0_, then later reverses its negative growth and returns to zero. If, as in Fig. 11, the pulse width is much larger than *τ*_*a*_, the step down response is simply the negative of the step-up response. For smaller step duration, the step-down response is still negative, but is not a mirror image of the step-up dynamics. Taking *s*_0_ to be positive, this antisymmetric response implies that as the eyespot rotates into the light there is a step-up response with *ω*_1_ > 0, corresponding to transient *cis* flagellar dominance, and when the eyespot rotates out of the light then *ω*_1_ < 0, associated with *trans* flagellum dominance. This is precisely the dynamics shown in Fig. 1 that allows monotonic turning toward the light as the cell body rotates.

Note that the adaptive dynamics coupling *ω*_1_, > *s*, and *h* is left unchanged by the simultaneous change of signs *ω*_1_ → −*ω*_1_, *h* → −*h*, and *s* → −*s*. This symmetry allows us to address positive and negative phototaxis in a single model, for if a step-up in light activates a transient dominant *trans* flagellum response in the cell orientation of Fig. 2, with *ω*_1_ < 0, we need only take *s* < 0.

Since the model (24) is constructed so that *ω*_1_ is forced by the signal *s*, the opposite-sign response to step-up and step-down signals is not an obvious feature. Yet, in the standard manner of coupled first-order ODEs, the hidden variable *h* can be eliminated, yielding a single, second-order equation for *ω*_1_. It can be cast in the simple form

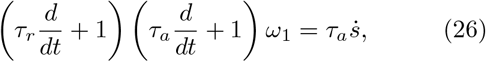

which is explicitly forced by the derivative of the signal, thus driven oppositely during and and down steps.

Previous studies [27] found a measurable time delay *τ*_*d*_ between the signal and the response that, in the language of the adaptive model, is additive with the intrinsic offset determined by the time scales *τ*_*r*_ and *τ*_*a*_. This can be captured by expressing the signal in (24) as *s*(*t* − *τ*_*d*_). The maximum amplitude of *ω*_1_(*t*) then occurs at the time

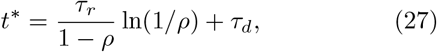

at which point the amplitude is *s*_0_*A*(*ρ*), where

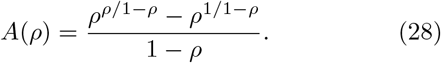

A fit to the step-response data yields *τ*_*d*_ = 28 ± 11 ms and *t*^*^ = 97 ± 18 ms. This delay between stimulus and maximum response has a geometric interpretation. The angle through which the eyespot rotates in the time *t*^*^ is |*ω*_3_|*t*^*^ = (0.28 ± 0.05)*π*, which is very nearly the angular shift *κ* = *π/*4 of the eyespot location from the (**ê**_2_-**ê**_3_) flagellar beat plane (Fig. 1). Since *ω*_3_ < 0, the eyespot *leads* the flagellar plane and thus *t*^*^ is the time needed for the beat plane to align with the light. In this configuration, rotations around **ê**_1_ are most effective [17, 27].

Since the function *A*(*ρ*) decreases monotonically from unity at *ρ* = 0, we identify the maximum angular speed 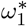 attainable for a given a stimulus as *s*_0_. With *s*_0_ = *gI*, we can remove *g* from the problem by instead viewing 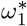 as the fundamental parameter, setting

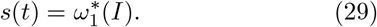

As indicated in (29), there is surely a dependence of 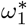 on the light intensity, not least because the rotational speed will have a clear upper bound associated with the limit in which the subdominant flagellum ceases beating completely during the transient photoresponse.

Turning now to the oscillating light signals experienced by freely-rotating cells, the directionality of the eyespot implies that the signal will be a *half-wave rectified* sinusoid (HWRS). Figure 11(b) shows the response of *ω*_1_ to two single half-period signals of this type. Compared to the square pulses of equal duration and maximum [Fig. 11(a)] the maximum response amplitude is reduced due to the lower mean value and slower rise of the signal. The frequency response of the adaptive model is most easily deduced from (26), and if 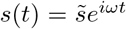, then there will be a proportionate amplitude 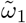. We define the response 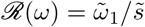, gain *G*(*ω*) = | *ℛ* (*ω*)| and phase shift *χ* = tan^−1^[Im(*ℛ*)*/*Re(*ℛ*)]. These are

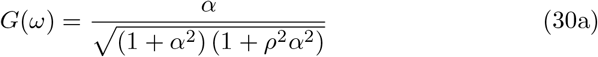

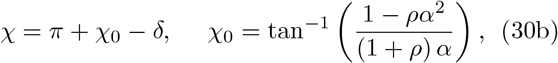

where *α* = *ωτ*_*a*_, *δ* = *ωτ*_*d*_ and the additive term of *π* in the phase represents the sign of the overall response. Figures 12(a,b) show these quantities as a function of the stimulus frequency *ω* for various values of *ρ*. The peak frequency is at 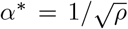, or 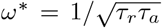, at which *G*(*ω*^*^) = 1*/*(1 + *ρ*) and *χ* = *π ω*^*^*τ*_*d*_. Fig. 12(a) shows that the peak is sharp for large *ρ* and becomes much broader as *ρ* → 0. The peak amplitude decays in a manner similar to (28) for a step response [Fig. 12(c)].

**FIG. 12.**
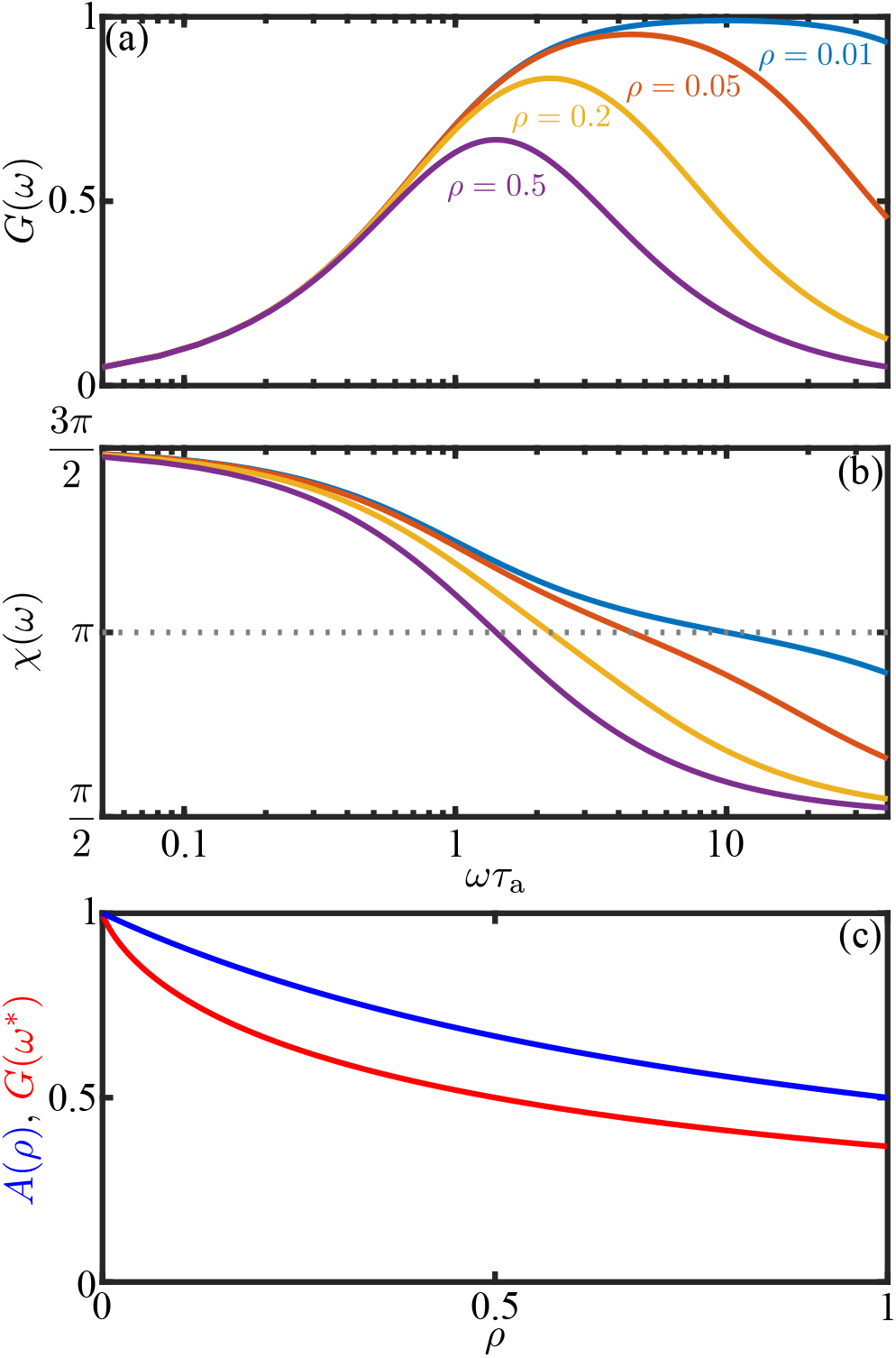
Dynamics of the adaptive model. (a) Gain (30a) for various values of *ρ* = *τ*_*r*_ */τ*_*a*_. (b) As in (a), but for the phase shift *χ* (30b), with *τ*_*d*_ = 0. (b) Comparison of peak amplitude for step-up and oscillatory forcing as a function of *ρ*.

The peaked response function amplitude (30a) and phase shift (30b) are qualitatively similar to those obtained experimentally by Josef, *et al*. [27], who analyzed separately the *cis* and *trans* responses and found distinct peak frequencies for the two, and investigated the applicability of more complex frequency-dependent response functions than those in (30). In the spirit of the analysis presented here we do not pursue such detailed descriptions of the flagellar responses, but it would be straight-forward to incorporate them as we discuss in Sec. IV B.

Using the same protocol as for the step function response in Fig. 10, we measured the frequency dependent photoresponse by subjecting cells to an oscillating light intensity at five distinct frequencies, analyzing the transient waveforms using RFT and determining the beat-average torque magnitude. The results of this study (Fig. 13), were fit to the form (30a), from which we obtained the time constants *τ*_*r*_ = 0.009 ± 0.002 s and *τ*_*a*_ = 0.52 ± 0.10 s, and thus *ρ* = 0.02. This strong separation between response and adaptation time scales is consistent with that seen under the step response (Fig. 10(a,b)) and leads to the broad peak of the frequency response curve. The peak frequency (≃2 Hz) is in very close agreement with recent direct measurements of rotation frequency about the cell body axis on free-swimming cells [18]. The phase data shown in Fig. 13(b) are well-described by the adaptive model with the parameters determined from the fit to the amplitude, with a time delay of *τ*_*d*_ = 38±5 ms, a value that is consistent with that obtained from the step response. Note that for frequencies *ω* near *ω*^*^ and for *ρ* ≪ 1, the phase has the simple form

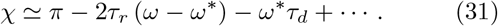

This result shows that while negative detuning from *ω*^*^ by itself increases the phase above *π*, the time delay can be a more significant contribution, leading to *χ* < *π*. Such is indeed the case in Fig. 13, where the peak frequency is ≃ 2 Hz, but |*ω*_3_|*τ*_*d*_ ≃ 0.13*π* and *χ*(*ω*^*^) < *π*.

**FIG. 13.**
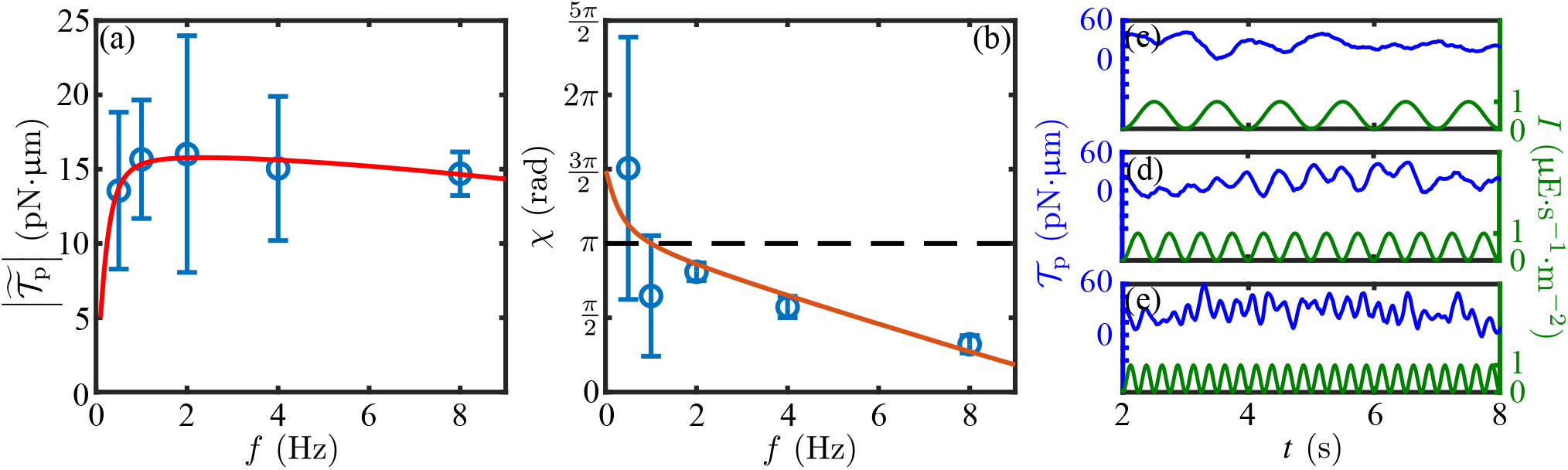
Frequency response of immobilized cells. (a) Measured beat-average phototactic torque determined from RFT (for positive phototaxis) at five stimulus frequencies (0.5, 1, 2, 4 and 8 Hz) for *n*_cells_ = 3 (blue) fitted to (30a) (red line). (b) As in (a), but for the phase *χ* of the response, fitted to (30b). The photoresponse shown (in blue) for three different stimulus frequencies (in green): (c) 1 Hz, (d) 2 Hz, and (e) 4 Hz.

## IV. DYNAMICS OF PHOTOTACTIC TURNS

### A. Helices, flagellar dominance, and eyespot shading

We now consider the larger length scales associated with the swimming trajectory of cells, and note the convention that rotation around an axis **e**_*i*_ is taken to have a *positive* angular velocity *ω*_*i*_ if the rotation is *clockwise* when viewed along the direction that **e**_*i*_ points. *Chlamydomonas* spins about **e**_3_ with an angular velocity *ω*_3_ < 0, and we define the positive frequency *f*_*r*_ = −*ω*_3_*/*2*π*. Its helical trajectories arise from an additional angular velocity 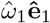, and we assume that *ω*_1_, *ω*_3_, and the translational speed *u* along **ê**_3_ are sufficient to define the trajectories, without invoking an angular velocity *ω*_2_.

The natural description of swimming trajectories is through the Euler angles (*ϕ, θ, ψ*) that define its orientation. In the standard convention [53], their time evolution is given by angular velocities (*ω*_1_, *ω*_2_, *ω*_3_) as follows,

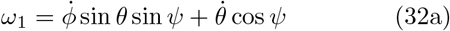

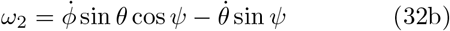

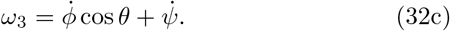

The transformation from the body frame **x** to the laboratory frame **x**^′^ is **x**^′^ = **A**·**x** and the reverse transformation is via **x** = **Ã** · **x**^′^, where **Ã** = **A**^−1^ = **A**^*T*^, with

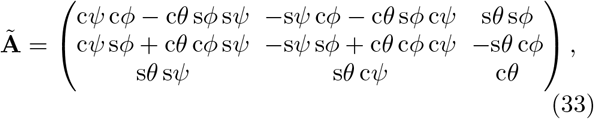

and we have adopted the shorthand *cψ* = cos*ψ*, etc.

The connection between helical swimming trajectories and the angular velocities *ω*_*i*_ has been made by Crenshaw [54–56] by first postulating helical motion and then finding consistent angular velocities. We use a more direct approach, starting from the Euler angle dynamics (32). If there is motion along a helix, and 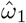 and *ω*_3_ are nonzero and constant, then apart from the degenerate case of orientation purely along **ê**_*z*_, where “gimbal locking” occurs, we must have 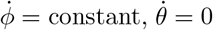 and 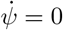. If we thus set 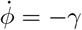 (the sign choice taken for later convenience), *θ* = *θ*_0_ (with −*π/*2 ≤ *θ*_0_ ≤ *π/*2), then a solution requires *ψ* = *π/*2 and the primary body axis is

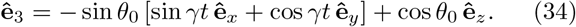

If the organism swims along the positive **ê**_3_ direction at speed *u*, then **ê**_3_ is the tangent vector 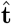 to its trajectory and we can integrate (34) using 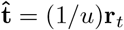 to obtain

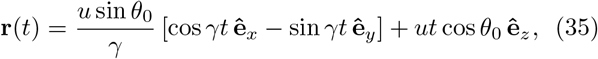

which is a helix of radius *R*_*h*_ and pitch *P*_*h*_ given by

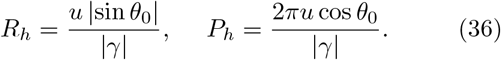

With the parameters (*γ, θ*_0_) taking either positive or negative values, there are four sign choices: (+, +), (+,−), (−, +), and (−, −). Since the *z* coordinate in the helices (35) increases independent of those signs, we see that when *γ* > 0 the *x* − *y* components of the helices are traversed in a clockwise (CW) manner and the helices are left-handed (LH), while when *γ* < 0 the in-plane motion is CCW, and the helices are right-handed.

We are now in a position to describe quantitatively the helical trajectories of swimming *Chlamydomonas* in the absence of photostimulation. From the estimated angular speed 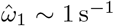 in (19), the typical value |*ω*_3_| ∼ 10 s^−1^ and swimming speed *u* ∼ 100 μm/s, we find *R*_*h*_ ∼ 1 μm and *P*_*h*_ ∼ 60 μm, both of which agree well with the classic study of swimming trajectories (Fig. 6 of [16]), which show the helical radius is a small fraction of the body diameter and the pitch is ∼ 6 diameters.

Next we describe in detail the helical trajectories adopted by cells in steady-state swimming, either toward the light during positive phototaxis, or away from it during negative phototaxis. For such motions, the relevant angular rotations are *ω*_3_ and the intrinsic speed 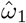. As noted earlier [17], there are four possible configurations to be considered on the basis of the sense of rotation around **ê**_1_ as determined by *cis* dominance 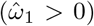 or *trans* dominance 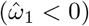. In both cases the relationship between the helix parameters *γ* and *θ*_0_ is

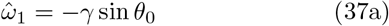

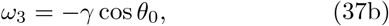

with *ω*_3_ < 0 in both cases. In *trans* dominance, 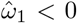 and a solution of (37) has 0 ≤ *θ*_0_ ≤ *π/*2, whereas for *cis* dominance, 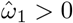 and −*π/*2 ≤ *θ*_0_ ≤ 0. Thus,

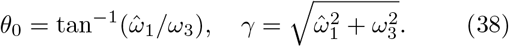

Setting 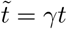 we obtain the helical trajectories

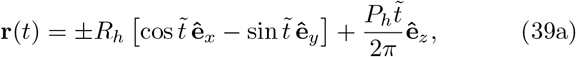

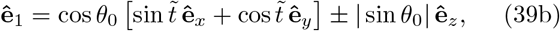

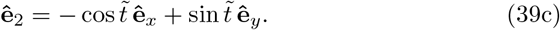

for *trans* (+) and *cis* (−) dominance.

We can now express quantitatively features regarding the eyespot orientation with respect to the helical trajectory that have been remarked on qualitatively [17]. While there is some variability in the eyespot location, it is typically in the equatorial plane defined by **ê**_1_ and **ê**_2_, approximately midway between the two. We take it to lie at an angle *κ* ∈ [0, *π/*2] with respect to **ê**_2_, such that the outward normal **ô** to the eyespot is

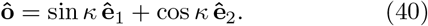

The outward normal vectors to the helix cylinder are 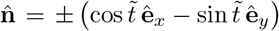, so the projection of the eyespot normal on 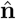 have the time-independent values

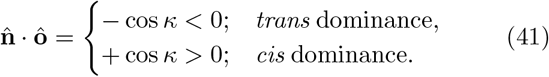

Thus, for any *κ* ∈ [0, *π/*2], the eyespot points to the *inside* (*outside*) of the helix for *trans* (*cis*) dominance. This confirms the general rule that any given body-fixed spot on a rigid body executing helical motion due to constant rotations about its axes has a time-independent orientation with respect to the helix. When *κ* = 0, **ô** = **ê**_2_, which points to the *cis* flagellum, and we see that the dominant flagellum is always on the *outside* of the helix.

If light shines long some direction 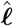, its projection on the eyespot is 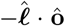; for light shining down the helical axis, 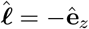, then the projections in the two cases are

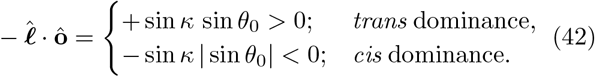

These various situations are summarized in Table II and shown in Fig. 14. As remarked many years ago, *trans* dominance holds in the case of negative phototaxis, and *cis* dominance in positive phototaxis. The conclusion from Table II is that when negatively phototactic cells swim away from the light or positively phototactic cells swim toward the light their eyespots are shaded, and thus the “stable” state is one minimal signal. so that when the first of these conditions holds during negative phototaxis, whereas the second occurs for positive phototaxis. Note that in the degenerate case *θ*_0_ = 0, when the helix reduces to a straight line, the projections (42) vanish, so a cell moving precisely opposite its desired direction would receive no signal to turn. A small helical component to the motion eliminates this singular case.

**TABLE II.** Left-handed helical swimming of *Chlamydomonas*.

| dominant flagellum | $\hat{\omega}_1$ | swimming relative to light source | eyespot orientation | eyespot status |
| --- | --- | --- | --- | --- |
| *cis* | + | toward | outside | shade |
| *cis* | + | away | outside | light |
| *trans* | − | toward | inside | light |
| *trans* | − | away | inside | shade |

**FIG. 14.**
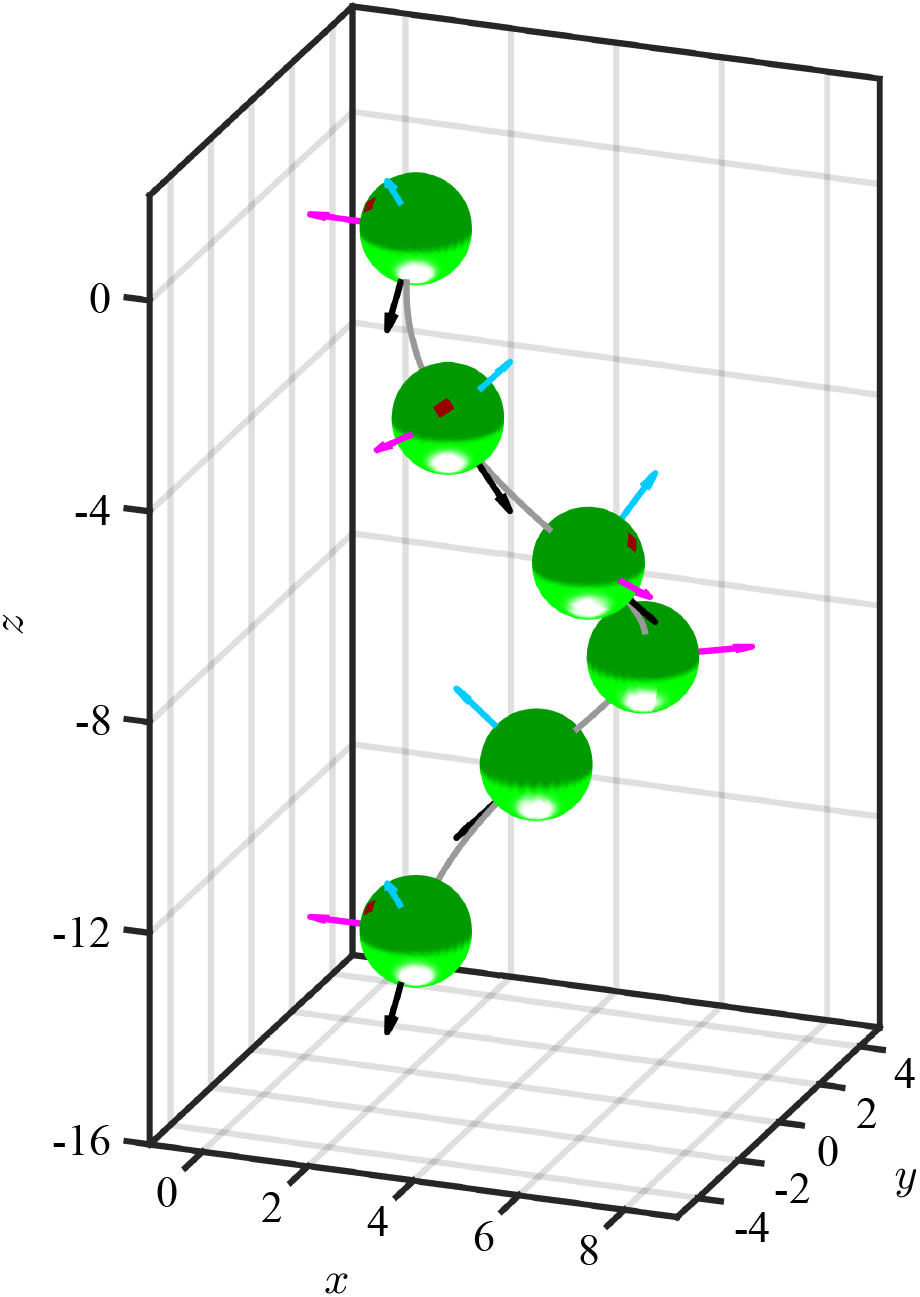
Helical trajectory of a *cis*-dominant cell which is swimming in a direction aligned with the light source located at the bottom. Principal body rotation axes are depicted as cyan, magenta and black arrows for **ê**_**1**_, **ê**_**2**_ and **ê**_**3**_ axes respectively. Eyespot is always located in the shading hemisphere of the cell body and is pointing outwards from the helix, as is the *cis* flagellum (not shown).

### B. Phototactic steering with adaptive dynamics

Now we merge the adaptive photoresponse dynamics with the kinematics of rigid body motion. The dynamics for the evolution of the Euler angles in the limit *ω*_2_ = 0 is obtained from (32), yielding

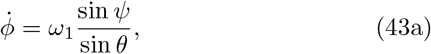

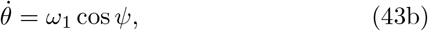

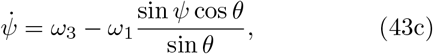

Given the assumption *ω*_2_ = 0, these are exact. As we take *ω*_3_ to be a constant associated with a given species of *Chlamydomonas*, it remains only to incorporate the dynamics of *ω*_1_ and the forward swimming speed to have a complete description of trajectories. The angular speed *ω*_1_ is the sum of intrinsic and phototactic contributions,

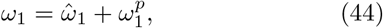

where 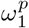 is described by the adaptive model.

It is natural to adopt rescalings based on the fundamental “clock” provided by the spinning of *Chlamydomonas* about **ê**_3_. Recalling that *ω*_3_ < 0, these are

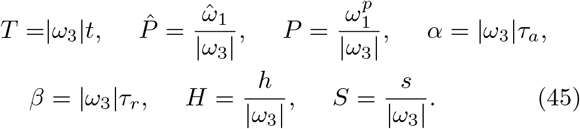

To incorporate the photoresponse, the light signal at the eyespot must be expressed in terms of the Euler angles. Henceforth we specialize to the case in which the a light source shines in the *x* − *y* plane along the negative *x*-axis, so 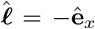, and the normalized projected light intensity 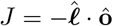 on the eyespot can be written as

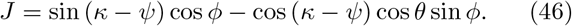

We assume for simplicity that eyespot shading is perfect, so that the signal sensed by the eyespot is

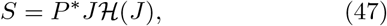

where 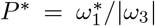 is negative (positive) for positive (negative) phototaxis, and ℋ is the Heaviside function.

With these rescalings, the dynamics reduces to

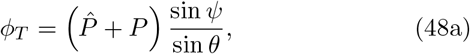

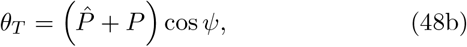

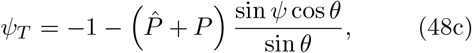

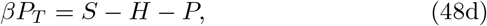

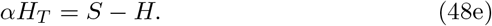

These five ODEs, supplemented with the signal definition in (46) and (47), constitute a closed system. To obtain the swimming trajectory, we use the cell body radius *R* to define the scaled position vector **R** = **r***/R*, so that the dynamics **r**_*t*_ = *u***ê**_3_ becomes

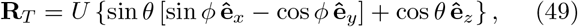

where *U* = *u/*(*R* |*ω*_3_|) is the scaled swimming speed. For typical parameter values (*u* = 100 μm/s, *R* ∼ 5 μm, and *f*_*r*_ = 1.6 Hz), we find *U* ∼ 2. Given (*θ*(*T*), *ϕ*(*T*)), we integrate (49) forward from some origin **R**(0) to obtain **R**(*T*), and use the triplet (*θ*(*T*), *ϕ*(*T*), *ψ*(*T*)) and the matrix **A**, the inverse of **Ã** in (33), to obtain **ê**_1_(*T*) and **ê**_2_(*T*).

An important structural feature of the dynamics is its partitioning into sub-dynamics for the Euler angles and the flagellar response. The connection between the two is provided by the response variable *P* (*T*), via the signal *S*, such that any other model for the response (for example, one incorporating distinct dynamics for the *cis* and *trans* flagella), or the signal (including only partial eyespot directionality, or cell body lensing) can be substituted for the adaptive dynamics with perfect shading.

The model (48) has 4 main parameters: 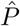 determines the unstimulated swimming helix, *P*^*^ sets the maximum photoresponse turn rate, and *α* and *β* describe the adaptive dynamics. Additional parameters are the eyespot angle *κ* (40) and time delay *τ*_*d*_. To gain insight, we first adopt the simplification that the eyespot vector **ô** is along **ê**_2_ (*κ* = 0), set *τ*_*d*_ = 0, and solve the initial value problem in which a cell starts swimming in the *x* − *y* plane (*θ*(0) = *π/*2) along the direction −**ê**_*y*_ (*ϕ*(0) = *π/*2) with its eyespot orthogonal to the light (*ψ*(0) = 0; **ê**_1_ = **ê**_*x*_ and **ê**_2_ = **ê**_*z*_) and about to rotate into the light. Figure 15 shows the results of numerical solution of the model for the non-helical case 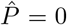, with *P*^*^ = − 0.4 for positive phototaxis, *α* = 7, *β* = 0.15 and *U* = 2. We see in Fig. 15(a) how the initially large photoresponse when the cell is orthogonal to the light decreases with each subsequent half turn as the angle *ϕ* evolves toward *π/*2 [Fig. 15(b)]. The signal at the eyespot [Fig. 15(c)], is a half-wave rectified sinusoid with an exponentially decreasing amplitude. Note that for this non-helical case the Euler angle *θ* remains very close to *π/*2 during the entire phototurn, indicating that the swimmer remains nearly in the *x* − *y* plane throughout the trajectory.

**FIG. 15.**
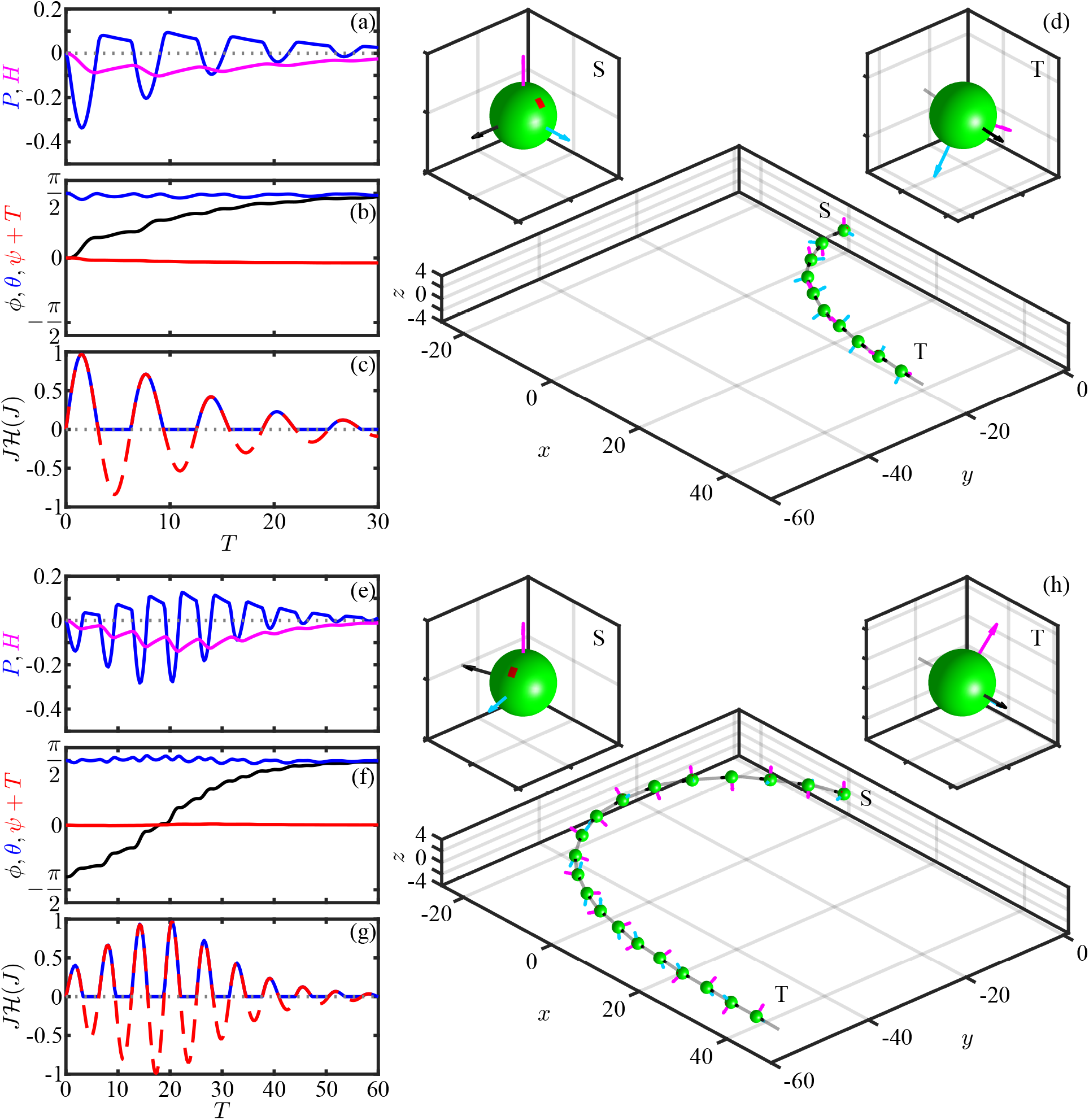
Positive phototaxis from model for non-helical swimming. (a-c) Time evolution of adaptive variables, Euler angles, and eyespot signal during a phototurn, i.e. from swimming orthogonal to the light to moving directly toward it. (d) Trajectory of the turn showing initial (S) and final (T) orientations of the cell (also in magnified insets). (e-h) Analogous to panels above but with dynamics starting from an orientation facing nearly away from the light. Parameters are 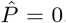, *P*^*^ = − 0.4, *α* = 7, *β* = 0.14, and *U* = 2, with the eyespot along **ê**_2_ (i.e. *κ* = 0 but shown in canonical position), and *τ*_*d*_ = 0. Color scheme of cell’s principal axes is same as in Fig. 14.

If the initial condition is nearly opposite to the light direction, the model shows that the cell can execute a complete phototurn (Fig. 15(e). In the language of Schaller, et al. [17], since the eyespot is “raked backward” when the cell swims toward the light, and the eyespot is shaded, when the cell swims away from the light it picks up a signal and can execute a full *π* turn to reach the light.

Next we include helicity in the base trajectory, setting 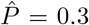. In the absence of phototactic stimulation this value leads to helical motion with a ratio of helix radius to pitch of 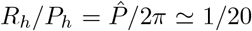, a value considerably larger than to that seen experimentally [16], but useful for the purposes of illustration. The phototurn dynamics shown in Fig. 16 exhibits the same qualitative features seen without helicity, albeit with much more pronounced oscillations in the evolution of the Euler angles, particularly of *ϕ* and *θ*. Averaged over the helical path the overall trajectory is similar to that without helicity, and does not deviate significantly from the *x* − *y* plane.

**FIG. 16.**
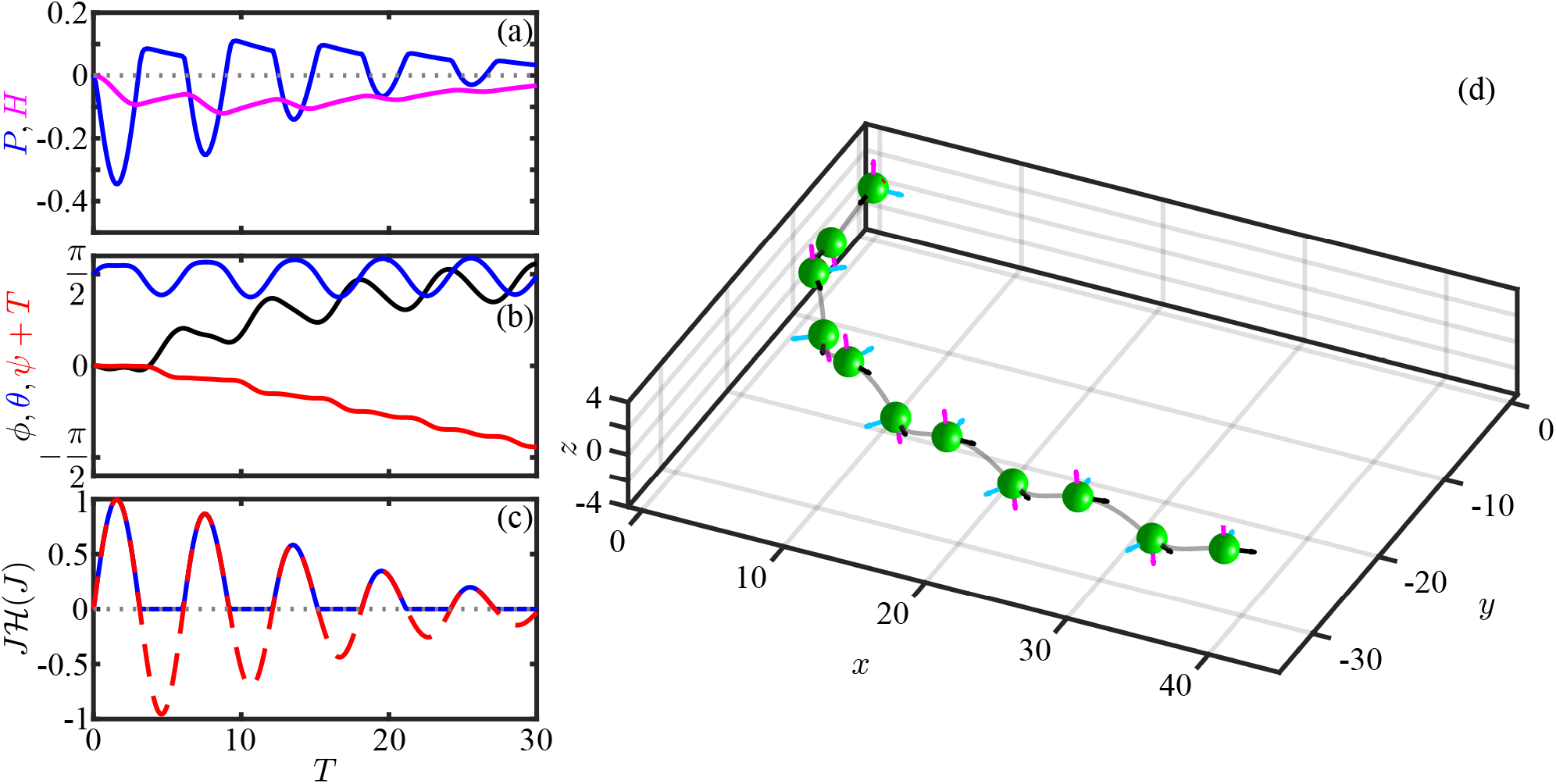
Positive phototaxis with helical swimming. As in Fig. 15 but with 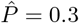.

To make analytical progress in quantifying a phototurn, we use the simplifications that are seen in Fig. 15 for the non-helical case. First, we neglect the small deviations of *θ* from *π/*2, and simply set *θ* = *π/*2. Second, we note that the time evolution of *ψ* is dominated by rotations around **ê**_3_, and thus we assume *ψ* = −*T*. This yields a simplified model in which the remaining Euler angle *ϕ* is driven by the cell spinning, subject to the adaptive dynamics.

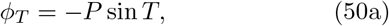

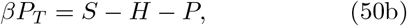

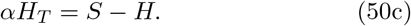

where we allow for a general eyespot location, using *J* = cos *ϕ* sin(*T* + *κ*), and thus

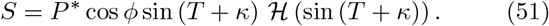

As it takes a number of half-periods of body rotation to execute a turn, we can consider the angle *ϕ* to be approximately constant during each half turn *n* (*n* = 1, 2, …) at the value we label *ϕ*_*n*_. For any fixed *ϕ*_*n*_, the signal *S* is simply a HWRS of amplitude *P*^*^ cos *ϕ*_*n*_. We explore two approaches to finding the evolution of *ϕ*_*n*_: (i) a *quasi-equilibrium* one in which the steady-state response of the adaptive system to an oscillating signal is used to estimate *P*, and (ii) a *non-equilibrium* one in which the response is the solution to an initial-value problem.

In the first approximation, we decompose the HWRS eyespot signal (51) into a Fourier series,

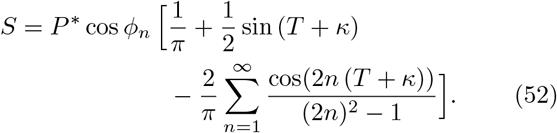

From the linearity of the adaptive model it follows that each term in this series produces an independent response with magnitude *G* and phase shift *χ* appropriate to its frequency *nω*_3_, for *n* = 0, 1, 2,.*…* Since the magnitude *G* in (30a) vanishes at zero frequency, the contributing terms in the photoresponse are

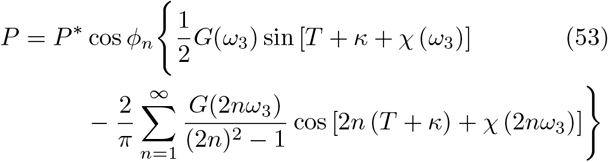

The first term dominates, as it is at the same frequency as the r.h.s. of the equation of motion *ϕ*_*T*_ = −*P* sin *T*. Keeping only this term, we integrate

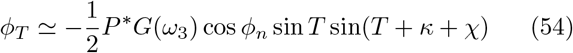

over one half period (*T* = *π*) and obtain the iterated map

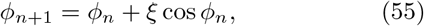

where

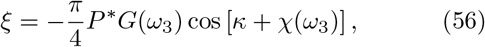

with *ξ* > 0 for *P*^*^ < 0 in positive phototaxis and *ξ* < 0 for negative phototaxis. An alternative approach involves the direct integration of the equations of motion over each half-turn. The lengthy algebra for this is given in Appendix A, where one finds a map analogous to (55), but with an *n*-dependent factor *ξ*_*n*_ that converges for large *n* to that in (56). Supplementary Video 2 [43] illustrates the cell reorientation dynamics under this map.

The iterated map (55) has fixed points at *ϕ*_±_ = ±*π/*2. Linearizing about those values by setting *ϕ*_*n*_ = ±*π/*2 + *δϕ*_*n*_, we obtain *δϕ*_*n*+1_ = (1∓*ξ*)*δϕ*_*n*_ and thus *δϕ*_*n*_ ∝ (1∓*ξ*)^*n*^*δϕ*_1_. Hence: (i) the angle +*π/*2 is stable for positive phototaxis when 0 ≤ *ξ* ≤2 and becomes unstable for *ξ* > 2, while it is unstable for negative phototaxis (*ξ* < 0). (ii) the angle −*π/*2 is unstable for positive phototaxis for any *ξ* > 0, while it is stable for negative phototaxis in the range −2 ≤ *ξ* ≤ 0 and unstable for *ξ* < −2. Thus, positively phototactic cells orient toward +*π/*2 and negatively phototactic cells orient toward −*π/*2, except for values of |*ξ*| > 2. These exceptional cases correspond to peak angular speeds 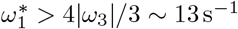.

Figure 17(a) shows the iterated map (55) for both positive and negative phototaxis. In the usual manner of interpreting such maps, the “cobwebbing” of successive iterations shows clearly how the orientation +*π/*2 is the global attractor for positive phototaxis, and *ϕ* = −*π/*2 is that for negative phototaxis. When |*ξ*| is small, the approach to the stable fixed point is exponential, *δϕ* ∼ exp(−*n/N*), where *N* = *N*_0_*/Q*(*ω*_3_), with

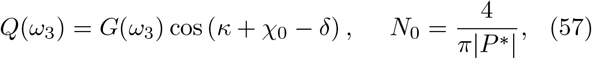

is the characteristic number half-turns needed for alignment. The number *N*_0_ reflects the bare scaling with the maximum rotation rate around **ê**_1_. For the typical value *P*^*^ ≃ 0.1 we have *N*_0_ ∼ 6. The presence of the gain *G* in denominator in (57) embodies the effect of tuning between the adaptation timescale and the rotation rate around **ê**_3_, while the term cos (*κ* + *χ*_0_ − *γ*) captures the feature discussed in frequency response studies in Sec. III C, namely that the flagellar asymmetries have maximum effect (and thus *Q* is maximized) when the negative phase shift *χ*_0_ − *δ* offsets the eyespot location). Figure 17(b) shows the dependence of *N/N*_0_ on the scaled relaxation time *f*_*r*_*τ*_*a*_ for various values of *ρ*. For the experimentally observed range *ρ* ≃ 0.02 − 0.1 there is a wide minimum of *N/N*_0_ around *f*_*r*_*τ*_*a*_ ∼ 1. This relationship confirms the role of tuning in the dynamics of phototaxis, but also shows the robustness of the processes involved.

**FIG. 17.**
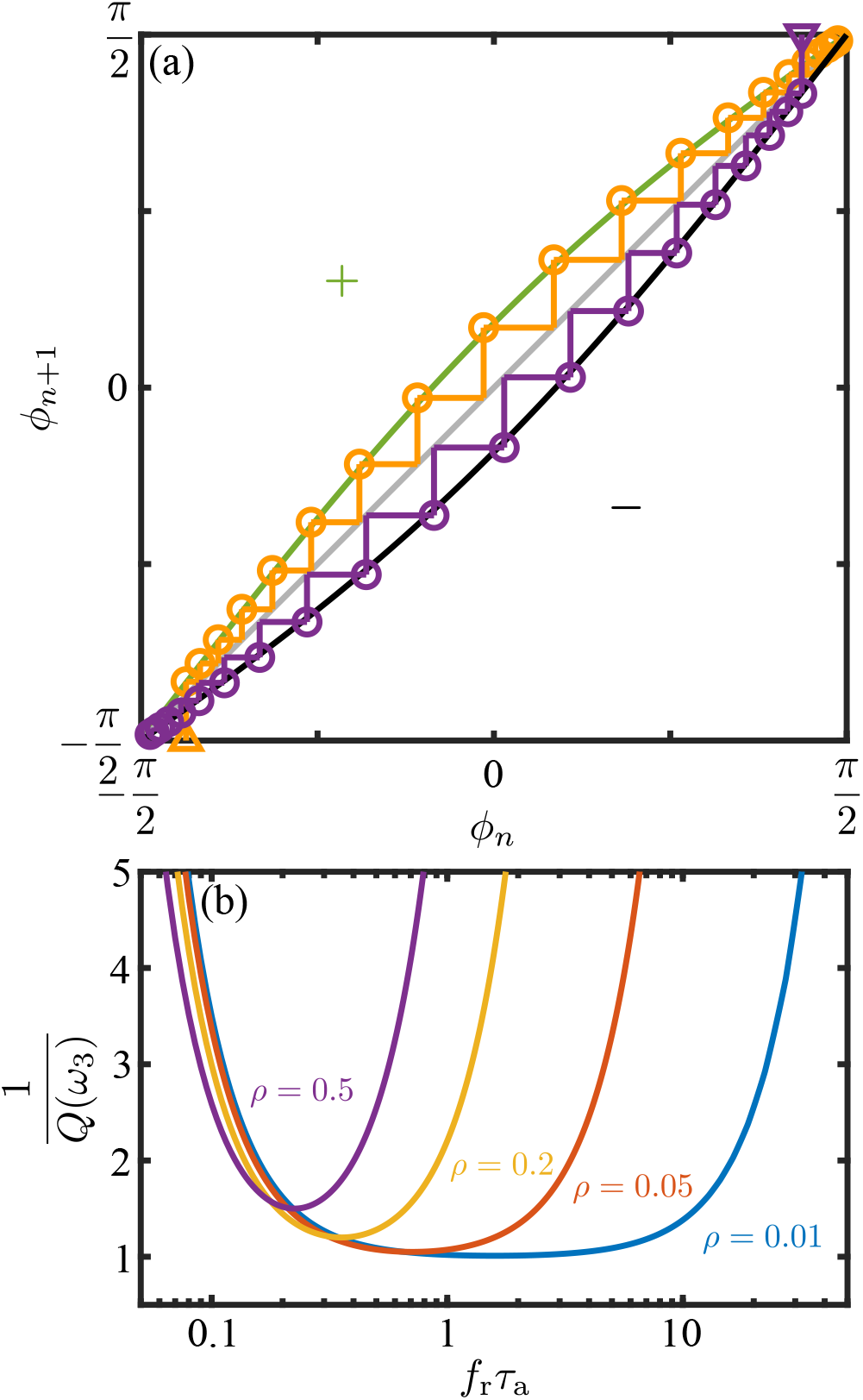
Iterated map of the reorientation model. (a) Cob-webbing of iterations starting from *ϕ* near − *π/*2 for positive phototaxis (upper branch in yellow) and near +*π/*2 for negative phototaxis (lower branch in purple), as indicated by triangles, for light shining toward −**ê**_*x*_. Values of *ξ* = ∓0.29 are used which are calculated based on (56) using values from Table III. (b) Response factor 1*/Q* in (57) for number of half-turns needed for alignment as a function of tuning parameter, for various values of *ρ* as indicated.

Returning to the evolution equation (54) for *ϕ*, we can also average the term sin *T* sin(*T* + *κ* + *χ*) over one complete cycle to obtain the approximate evolution equation

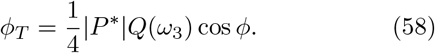

For positive phototaxis and with *ϕ*(0) = 0, the solution to this ODE can be expressed in unrescaled units as

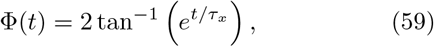

where Φ = *π/*2 + *ϕ*, with Φ(0) = *π/*2, Φ(∞) = *π*, and the characteristic time in physical units is

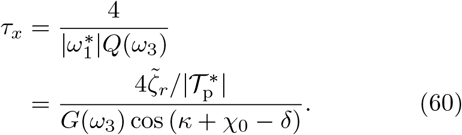

This is a central result of our analysis, in that it relates the time scale for reorientation during a phototurn to the magnitude and dynamics of the transient flagellar asymmetries during the photoresponse. As discussed above, the function *Q*(*ω*_3_) embodies the optimality of the response—in terms of the tuning between the rotational frequency and the adaptation time, and the phase delay and eyespot position—but also captures the robustness of the response through the broad minimum in *Q* as a function of both frequency and eyespot position.

Using the 3D tracking system described in Sec. II, we analyzed 6 pairs of movies, within which we tracked 283 trajectories with duration greater than 10 s and which included the trigger frame. From those, 44 showed both positive phototaxis and included a full turn to *ϕ* = *π/*2 as shown in Fig. 18(a) and Supplementary Video 3 [43]. These trajectories were cropped to include any points for which − *π/*2 ⩽ *ϕ* ⩽ *π/*2 and which could then be fitted to (59) to determine the experimental time constant *τ*_*x*_. The boxplot in Fig. 18 shows the experimental mean value *τ*_*x*_ = 0.96 ± 0.36 s (also in Table III). This can be compared to the estimates obtained within the steady-state approximation (60) and the transient analysis (Appendices A and B), both of which are based on the mean and standard error value of the peak flagellar phototorque of −35.0±8.8 pN·μm, the mean values of the adaptive flagellar response time-scales (*τ*_*r*_ = 0.018 s and *τ*_*a*_ = 0.764 s in Table III) and the effective drag coefficient 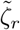. The steady-state estimate of *τ*_*x*_ is 1.04±0.26 s (with *Q/*4 = 0.243), while the transient estimate is 0.96±0.24 s (with the nonequilibrium counterpart 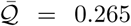 in (B5)). This agreement provides strong validation of the model of adaptive phototaxis developed here.

**FIG. 18.**
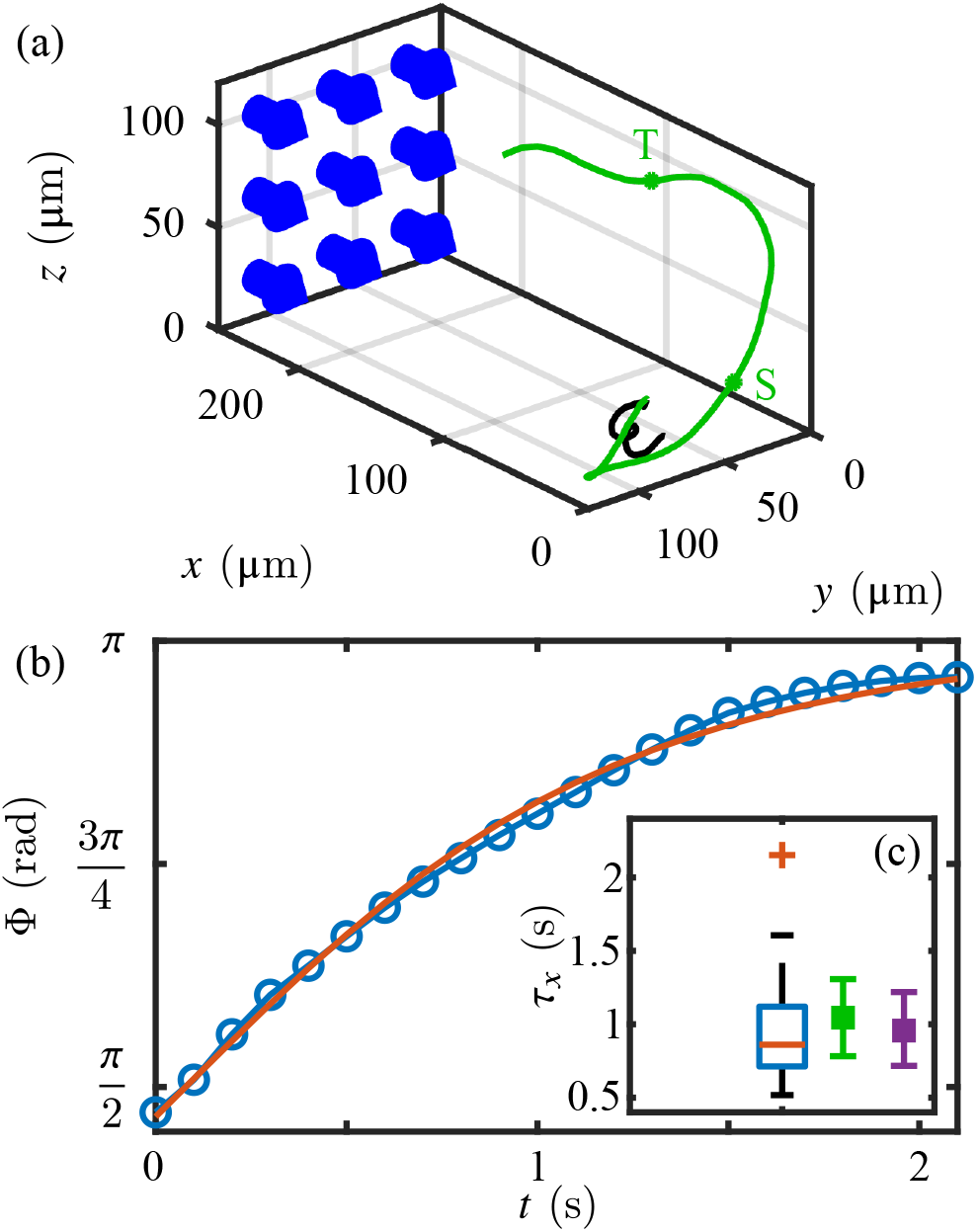
Phototactic swimmers tracked in three-dimensions. (a) A U-turn: trajectory in black is prior to light stimulation, that in green is afterwords. Blue arrows indicate direction of light. The cropped trajectory used for fitting the reorientation dynamics is bounded by the points S and T. (b) Dynamics of the reorientation angle Φ (blue) for the cropped trajectory fitted using (59). (c) Box plot of distribution of fitted *τ*_*x*_ with estimates, steady-state (green) and nonequilibrium (purple), derived from micropipette experiments.

**TABLE III.** Time scales that define the reorientation dynamics. Experimental data are from the present study except for the cell body rotation frequency.

| Quantity | Symbol | Mean ± SD |
| --- | --- | --- |
| adaptation time | $\tau_a$ | $0.764 \pm 0.190$ s |
| response time | $\tau_r$ | $0.018 \pm 0.009$ s |
| time delay | $\tau_d$ | $0.028 \pm 0.011$ s |
| cell body rotation frequency [18] | $f_r$ | $1.67 \pm 0.35$ Hz |
| reorientation time scale | $\tau_x$ | $0.96 \pm 0.36$ s |

## V. DISCUSSION

This study has achieved three goals: the development of methods to capture flagellar photoresponses at high spatio-temporal resolution, the estimate of torques generated during these responses and the measuremnt of relevant biochemical time scales that underlie phototaxis, and the integration of this information into a mathematical model to describe accurately the phototactic turning of *Chlamydomonas*. In developing a theory for phototurns, our work also puts on a more systematic mathematical foundation qualitative arguments [17] for the stability of phototactic trajectories based on eyespot orientation in both positive and negative phototaxis.

We have emphasized that rather than seek to develop a maximally detailed model of the dynamics of individual flagellar responses involved in phototaxis, we aimed to provide, in the context of one simple microscopic model, a multiscale analysis of the connection between such responses and the phototactic trajectories in a manner than can be easily generalized. Thus we obtain from experiment the values for microscopic and macroscopic time scales, as shown in Table III, and derive relation between them, culminating in (60) (and (B5)).

This analysis highlights the dual issues of optimatility and robustness. As noted in the introduction, the former was first addressed using a paralyzed-flagella mutant strain (*pf14*) and an electrophysiological approach on a bulk sample by [25]. In those experiments, a suspension of immotile cells was exposed to an oscillating light stimulus (wavelength 500 nm) and the resulting photoreceptor current was measured in a cuvette attached to two platinum electrodes. The experiment using relatively high light intensities observed a frequency response peak of 1.6 Hz when stimulated with ≈ 160 μE·s^−1^·m^−2^ and a frequency response peak of 3.1 Hz when stimulated with ≈ 40 μE·s^−1^·m^−2^. The former observation is in very good agreement with our results in Fig. 13 (peak response at ≃2 Hz), even though we used light stimulus intensities of ≈1 μE·s^−1^·m^−2^. We have not seen any evidence of cells having flagellar photoresponse dynamics that would corroborate the latter result of 3.1 Hz and this is a matter open to further study.

In addition, this study has addressed issues relating to past observations. With respect to the lag time *τ*_*d*_ of the photoresponse, we have measured by detailed study of the flagellar waveforms a value of 28±11 ms that is very similar to the value 30−40 ms observed earlier [21]. In addition, we have shown through the adaptive dynamics that the peak flagellar response is at a larger total delay time *t*^*^ given by (27) that corresponds accurately to the time between the eyespot receiving a light signal and the alignment of the flagellar beat plane with the light. Analysis of the phototactic model reveals that such tuning shortens the time for phototactic alignment.

Regarding the amount of light necessary for a flagellar photoresponse appropriate to positive phototaxis, we have converged, through trial and error, to ≈ 1 μE·s^−1^·m^−2^ at a wavelength of 470 nm. While this value is much lower than in other photoresponse experiments [26] where ≈60 μE·s^−1^·m^−2^ were used at a longer wavelength (543 nm), it is consistent with the sensitivity profile of channelrhodopsin-2 [57]. More detailed studies of the wavelength sensitivity of the flagellar photoresponse should be carried out in order to reveal any possible wavelength dependencies of quantities such as the time constants *τ*_r_ and *τ*_a_. Our work has addressed the relationship between the stimulus and the photoresponse of *Chlamydomonas* using an adaptive model that has perhaps the minimum number of parameters appropriate to the problem, each corresponding to a physical process. Attempts to derive similar relationships between stimulus and photoresponse [27] used linear system analysis. The result of such a signal-processing oriented method usually includes a much larger number of parameters necessary for the description of the system, without necessarily corresponding to any obvious measurable physical quantities. The evolutionary perspective that we emphasized in the introduction, culminating in the results presented in Fig. 2, points to several areas for future work. Chief among them is an understanding of the biochemical origin of the response and adaptive timescales of the photoresponse, in light of genomic information available on the various species. Flagellar and phototaxis mutants will likely be important in unravelling whether these time scales are associated with the axoneme directly or arise from coupling to cytoplasmic components. Additionally, we anticipate that directed evolution experiments such as those already applied to *Chlamydomonas* [58] can yield important information on the dynamics of phototaxis. For example, is it possible to evolve cells that exhibit faster phototaxis, and if so, which aspect of the light response changes? For the multicellular green algae, these kinds of experiments may also impact on the organization of somatic cells within the extracellular matrix, which has been shown to exhibit significant variability [59].

Another aspect for future investigation sits within the general are of control theory; the adaptive phototaxis mechanism that is common to the Volvocine algae, and to other systems such as *Euglena* [60], is one in which a chemomechanical system achieves a fixed point by evolving in time so as to null out a periodic signal. Two natural questions arise from this observation. First, what evolutionary pathways may have led to this behavior? Second, are there lessons for control theory in general and perhaps even for autonomous vehicles in particular that can be deduced from this navigational strategy?

We close by emphasizing that the flagellar photoresponse – and by extension phototaxis – is a complex biological process encompassing many variables, and that in addition to the short-term responses to light stimulation studied here there are issues of long-term adaptation to darkness or phototactic light that have only recently have begun to be addressed [61]. Together with the dynamics of phototaxis in concentrated suspensions [62], these are important issues for further work.

## ACKNOWLEDGMENTS

We thank Pierre A. Haas and Eric Lauga for very useful discussions and a critical reading of the manuscript, Kirsty Y. Wan for sharing code from previous work on flagellar tracking, David-Page Croft, Colin Hitch, John Milton, and Paul Mitton or technical support, and Ali Ghareeb for assistance with the fiber coupling apparatus. This work was supported in part by Wellcome Grant 207510/Z/17/Z. For the purpose of open access, the authors have applied a CC BY public copyright license to any Author Accepted Manuscript version arising from this submission. Additional support was provided by Grant No. 7523 from the Marine Microbiology Initiative of the Gordon and Betty Moore Foundation, and EPSRC Established Career Fellowship EP/M017982/1.

## Appendix A: Details of the iterated map from solution of the initial value problem

Here we provide details of the derivation of the iterated map for phototurns based on explicit solution of the initial value problem for the adaptive response. For conciseness we fix the eyespot position at *κ* = 0 and set the time delay

*τ*_*d*_ = 0. We start from the dynamics (50), rewritten for each full turn *n* ≥ 0 as

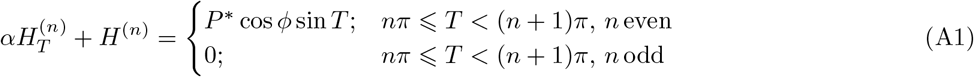

and

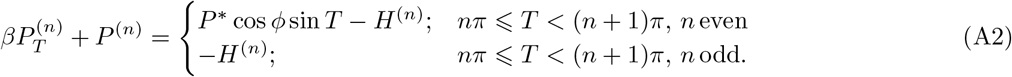

Solving this in a piecewise fashion we obtain

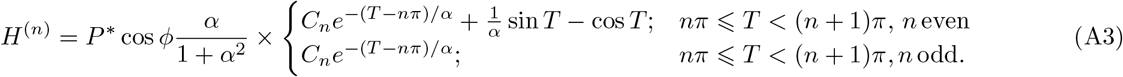

where *C*_*n*_ = (1− *r*^*n*+1^)*/*(1− *r*) and *r* = *e*^−*π/α*^. Continuity of *H* at the end of each light interval can be verified by noting that *H*^(*n*)^(*T* = (*n* + 1)*π*) ∝ *rC*_*n*_ + 1 for even *n*, while *H*^(*n*)^(*T* = *nπ*) ∝ *C*_*n*+1_ for the subsequent odd *n*, and observing that 1 + *rC*_*n*_ = *C*_*n*+1_.

The solution for the photoresponse variable can be expressed as 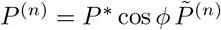, where

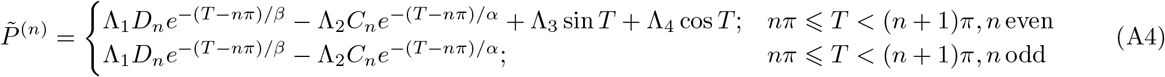

with *D*_*n*_ = (1 − *q*^*n*+1^)*/*(1 − *q*), *q* = *e*^−*π/β*^, and

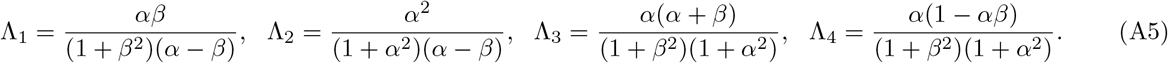

Since *n* represents the number of half-turns, with even(odd) values for the illuminated(shaded) periods, we integrate for each value of *n* ⩾ 0 to obtain

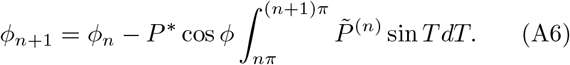

This has the form of (55), but with an *n*-dependent *ξ*_*n*_,

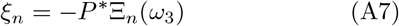

where

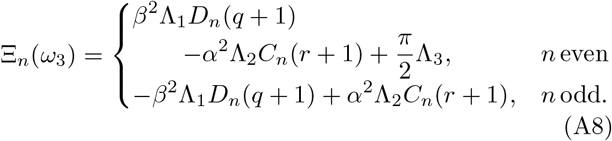

From the general structure of the iterated map, it is clear that the larger is *ξ*_*n*_ the larger the angular change within a given half-turn. It is of interest then to consider the average 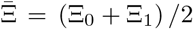 over the first two half turns, which gives the average coefficient

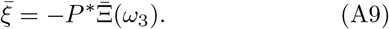

The quantity 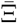 can be interpreted as the initial photoresponse function analogous to the steady-state response embodied in the amplitude *G*(*ω*_3_) and phase *χ*_0_ in (30). The functions *G*(*ω*_3_) cos *χ*_0_(*ω*_3_) and 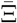 are compared in Fig. 19, where we see that the transient response function 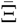 is about 10% higher at its peak, a feature that can be attributed to the fact that the hidden variable *H* has not yet built up to its steady value. But the two functions are otherwise remarkably similar, indicating the accuracy of the steady-state approximation.

**FIG. 19.**
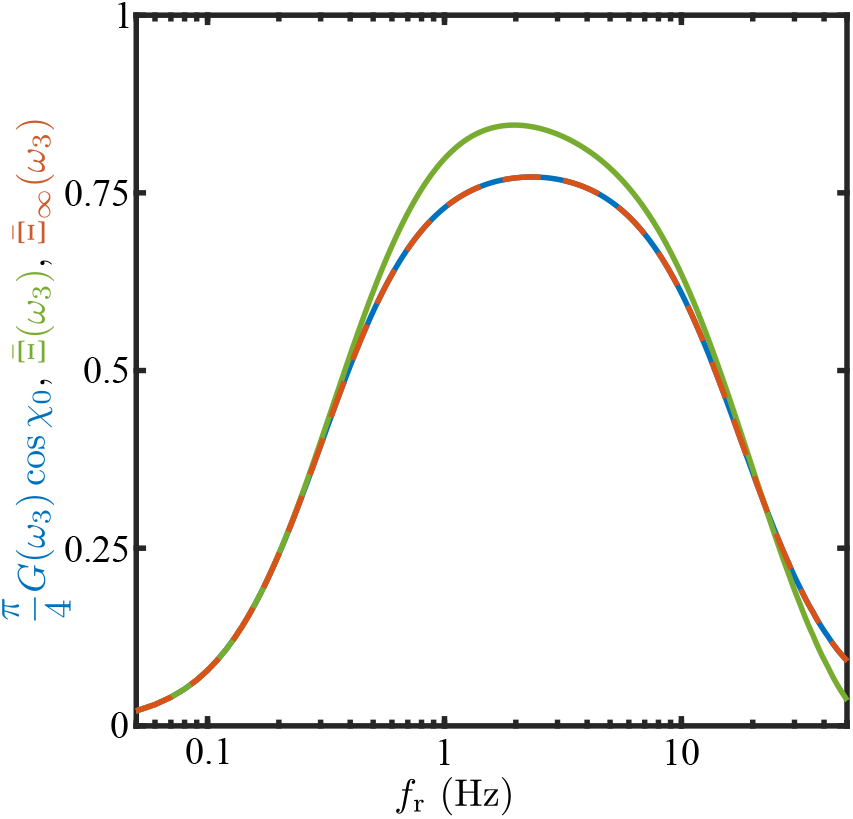
Response functions. For *κ* = *τ*_*d*_ = 0, the graph compares the steady-state response function (*π/*4)*G*(*ω*_3_) cos *χ*_0_ in (30) (dashed blue), the initial average response 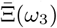 in (A9) (green) and the *n*→ ∞ limit of the transient response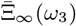 (dashed red), as functions of *f*_*r*_ = *ω*_3_*/*2*π*.

More generally, the coefficients Ξ_*n*_ exhibit an oscillating decay with *n*, converging as *n* → ∞ to

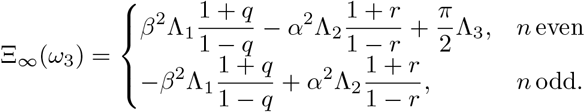

The connection to the steady-state approximation is obtained by considering the average 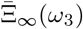 over the light and dark cycles (hence over even and odd values of *n*), where one finds 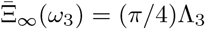, or

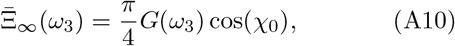

completely consistent with the steady-state analysis (56).

In the animation shown in Supplementary Video 2 [43] of the cell reorientation dynamics we evolve the iterated map using the {*ξ*_*n*_} and linearly interpolate between the values *ϕ*_*n*_ to obtain a smooth function of time.

## Appendix B: Details of continuous model

Using the solution of the initial value problem we can compute the continuous approximation to the evolution equation for *ϕ* by integrating over the fast photoresponse variables within a turn. From the governing equation 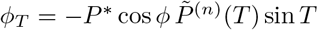, we obtain

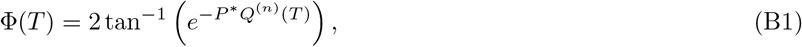

where

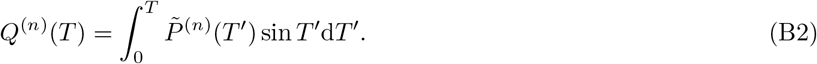

From (A4) we find

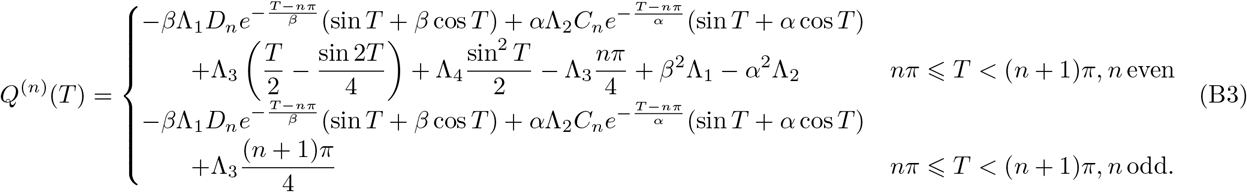

As shown in Figure 20, the function *Q*^(*n*)^(*T*) typically increases monotonically with *T*, exhibiting small oscillations around an interpolant that grows nearly linearly with time. These magnitudes of these oscillations vary between the light and dark halves of each turn. To quantify this asymmetry we compute the values *Q*^(*n*)^(*nπ*) at the start of each half turn,

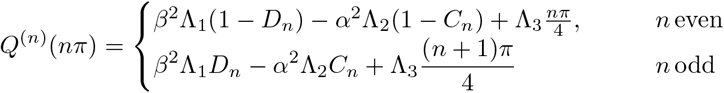

and the gradients 𝒬_*n*_ = [*Q*^(*n*+1)^((*n* + 1)*π*) − *Q*^(*n*)^(*nπ*)]*/π* of line segments connecting the node. One can easily show that 𝒬_*n*_ = Ξ_*n*_*/π*. The light-dark variation of these slopes serves as a measure of the smoothness of the reorientation dynamics, and from the first two values 𝒬_0_ and 𝒬_1_ we define two relevant quantities: the *strength* of the initial response as measured by the average of the slopes of the first two line segments 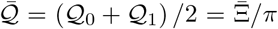, and its *smoothness*, as measured by the ratio 𝒴 = 𝒬_1_*/*𝒬 _0_.

**FIG. 20.**
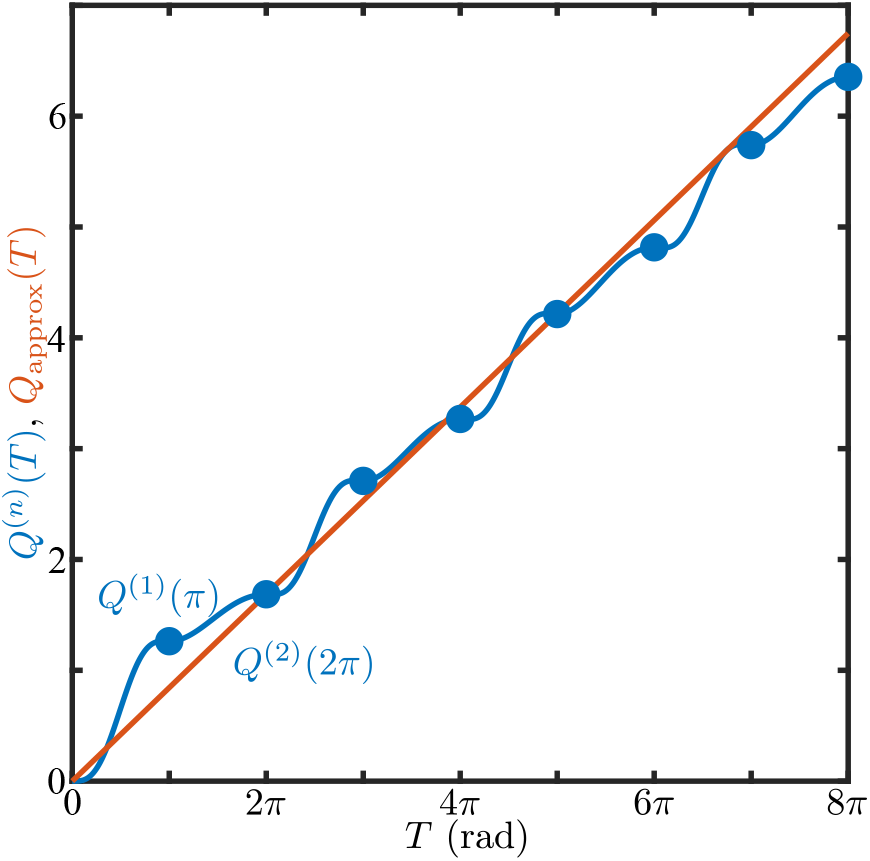
The functions *Q*^(*n*)^(*T*). Approximating *Q*^(*n*)^(*T*) as a straight line 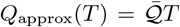. Both *Q*^(*n*)^(*t*) and 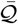 were computed with the experimentally derived *τ*_*r*_ = 0.009 s and *τ*_*a*_ = 0.520 s. The value of *f*_*r*_ was taken to be 1.67 Hz. The value of 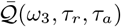 was calculated to be 0.265.

If, as in Fig. 20, we approximate *Q*^(*n*)^(*T*) by the line 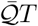, then the reorientation dynamics (B2) takes the simple form

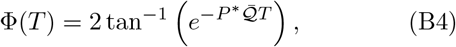

from which we identify the characteristic relaxation tim *τ*_*x*_ (in physical units) analogous to (60),

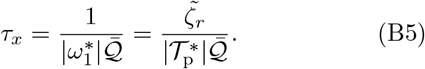

Finally, we explore the space of reorientation dynamics by probing *Q*^(*n*)^(*T*) through its dependency on parameters *α* and *β*. Our strategy is to observe how the quantities 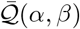 (Fig. 21(a)) and 𝒴 (*α, β*) (Fig. 21(b)), which are also functions of *α* and *β*, and essentially describe the curve’s shape, vary. Firstly, we make the observation that the (*α,β*) pairs acquired from micropipette experiments (step-up and frequency response; Fig. 10(b)) lie in the high-slope area 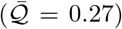 of the 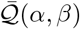 function (Fig. 21(a) and Fig. 21(c)). We also observe that the same data lie in an area of relatively moderate symmetry (𝒴 ≈ 0.34) of the (*α, β*) function (red markers in Fig. 21(b)), as opposed to the extreme cases of highest symmetry i.e. 𝒴 ≈ 0.7 (solid purple square in Fig. 21(b) and Fig. 21(d)) and lowest symmetry i.e. 𝒴 ≈ 0 (solid purple triangle in Fig. 21(b) and Fig. 21(e)).

**FIG. 21.**
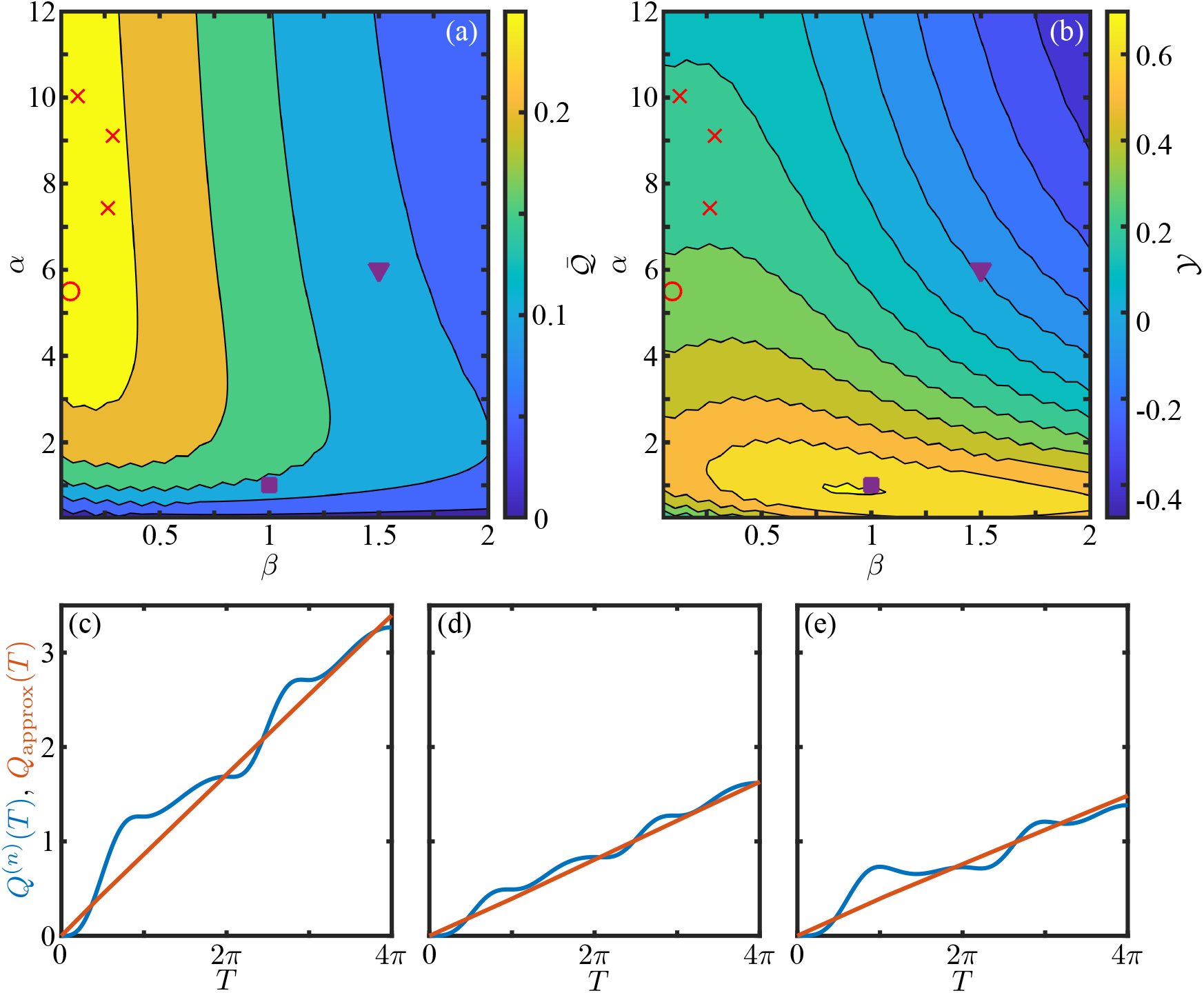
Contour plot maps of 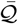 (a) and 𝒴 (b) with (*α,β*) pairs acquired from micropipette experiments shown with red markers (step-up as “x” and frequency as “o”). (c-e) Plots of *Q*^(*n*)^(*T*) (blue line) and linear approximation 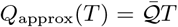 (red line) for three (*α,β*) value pairs shown in (a) and (b) as circle, square and triangle respectively. (c) is based on experimental data (open red circle), (d) corresponds to 𝒴 ≈ 0.7 (solid purple square) and (e) corresponds to 𝒴 ≈ 0 (solid purple triangle).

